# Diversification of a cryptic radiation, a closer look at Madagascar’s recently recognized bird family

**DOI:** 10.1101/825687

**Authors:** Jane L. Younger, Nicholas L. Block, Marie J. Raherilalao, J. Dylan Maddox, Kristen S. Wacker, Christopher C. Kyriazis, Steven M. Goodman, Sushma Reddy

**Affiliations:** Milner Centre for Evolution, Department of Biology & Biochemistry, University of Bath, Claverton Down, Bath, BA2 7AY, UK.; Department of Biology, Loyola University Chicago, 1032 W. Sheridan Road, Chicago, IL 60660, USA; Biology Department, Stonehill College, 320 Washington Street, Easton, MA 02357, USA; Association Vahatra, BP 3972, Antananarivo 101, Madagascar; Mention Zoology and Animal Biodiversity, University of Antananarivo, BP 906, Antananarivo (101), Madagascar; Field Museum of Natural History, 1400 S. Lake Shore Drive, Chicago, IL 60605, USA; Environmental Sciences, American Public University System, Charles Town, WV 25414, USA; Department of Ecology and Evolutionary Biology, University of Michigan, 1105 North University Avenue, Ann Arbor, MI 48109, USA; Department of Ecology and Evolutionary Biology, University of California Los Angeles, 612 Charles E. Young Drive South, Los Angeles, CA 90095, USA; Bell Museum of Natural History and Department of Fisheries, Wildlife, and Conservation Biology, University of Minnesota, St. Paul, MN 55108, USA

**Keywords:** Passeriformes, Bernieridae, tetrakas, phylogenetics, UCEs

## Abstract

Despite its status as a global biodiversity hotspot there is still much to be discovered about the birds of Madagascar, including a full accounting of species-level diversity and the avifauna’s origins. The Bernieridae is a Malagasy endemic family that went unrecognized by science for decades and unnamed until 2010. This cryptic family has long represented a missing piece of the puzzle of the avian tree of life. We present the first comprehensive phylogeny of Bernieridae in order to examine its diversification history on Madagascar and its place within Passeriformes. In light of recent discoveries of cryptic species-level diversity in Madagascar’s vertebrate fauna, we used broad geographic sampling and integrative taxonomic methods to investigate the potential for cryptic lineages within every known species of the Bernieridae. Our approach combines phylogenomics using ∼4500 loci of ultraconserved elements (UCEs), genetic clustering of thousands of single nucleotide polymorphisms (SNPs), and statistical analysis of morphological variation. These methods yielded the discovery of two unrecognized species in the previously monotypic genus *Bernieria*, along with new insights into patterns of fine-scale endemism in Madagascar’s humid forests. Our phylogenomic analyses provide conclusive support for Donacobiidae and Bernieridae as sister families, a biogeographically intriguing result given that the former is restricted to the Neotropics. We found a significant decline in the rate of speciation over time on Madagascar, consistent with a model of adaptive radiation. Bernieridae therefore joins the Vangidae as a second avian adaptive radiation on the island of Madagascar. These insights into the evolution of Bernieridae represent a step forward in understanding the origins and diversity of Madagascar’s endemic avifauna.

## Introduction

Islands are natural laboratories for studying the evolutionary processes of how species colonize and diversify in novel environments (Losos and Ricklefs 2009; Whittaker et al. 2017). Founding lineages can have varied responses to their new environments, from persisting with little change, to diversifying into adaptive radiations. Such island radiations often exhibit high phenotypic diversity, providing dramatic examples of the potential for rapid evolutionary change in response to new ecological interactions and habitats. Recognizing these endemic radiations has sometimes been difficult because of their extreme morphological disparity (Givnish et al. 2008; Reddy et al. 2012; Seehausen 2006). On highly isolated islands, classic examples of adaptive radiation, such as Darwin’s Finches and Hawaiian Honeycreepers, are still recognizable as cohesive groups due to the lack of comparable taxa nearby. However, distinguishing endemic radiations from a suite of independent colonization events can be more challenging in island systems with morphologically similar species on neighboring source continents.

In birds, molecular techniques have been key to uncovering relationships among disparate groups, leading to many rearrangements and reclassifications across the bird tree of life (Barker et al. 2004; Hackett et al. 2008). One of the most spectacular was the discovery of an unrecognized family endemic to Madagascar, Bernieridae, comprised of species previously placed in several Passeriformes families (Cibois et al. 1999; Cibois et al. 2001). The cryptic family was officially named in 2010 (Cibois et al. 2010). This discovery was only possible with genetic data and makes clear the need for robust phylogenies for examining the evolution of potentially convergent traits in ecology, behavior, biogeography, and diversification studies.

Madagascar is the 4^th^ largest island in the world and has been isolated from other landmasses since the breakup of Gondwana. The modern Malagasy avifauna, like most of the island’s terrestrial fauna, is the result of dispersals during the Cenozoic (Samonds et al. 2013; Vences et al. 2009; Yoder and Nowak 2006). Despite its position only 400 km to the east of Africa, relatively few African lineages have managed to colonize the island, possibly due to westerly winds and ocean currents (Ali and Huber 2010). Although species poor for its size (Schulenberg 2003), Madagascar has representatives of numerous lineages from across the bird tree of life. Many endemic species were previously thought to be single species dispersals of larger continental families (Schulenberg 2003). However, more recent analyses have shown several of these taxa to belong to endemic radiations, and their resemblance to African or Asian groups to be convergent (Cibois et al. 2001; Johansson et al. 2008b; Reddy et al. 2012). It is now clear that the Malagasy avifauna is more distinct than previously assumed and many of the widespread continental groups found on Asia and Africa do not have Malagasy representatives.

There is still much that is unknown about Madagascar’s unique biodiversity, including the presence of cryptic or hidden species. Recent phylogeographic studies have uncovered cryptic differentiation across populations restricted to different habitats across the island, including in mouse lemurs (Weisrock et al. 2010), tenrecs (Everson et al. 2018), frogs (Brown et al. 2014), and birds (Younger et al. 2018). In birds, well-sampled genetic studies based on broad geographical field surveys have led to the description of several new species in the last decade (Goodman et al. 2011; Younger et al. 2019; Younger et al. 2018). These studies indicate that more analyses are needed to gain a complete picture of the true diversity across this large and heterogenous island. A full accounting of species level diversity is crucial for both understanding evolutionary processes and effective conservation programs.

Little is known about the ecology and behavior of the Bernieridae, commonly known as tetrakas or Malagasy warblers. All species glean insects and are restricted to forests, with eight of the currently 11 species found in the Eastern Humid Forest. All members of this family were previously classified in different passerine families based on superficial similarities, mainly size and coloration. *Xanthomixis* and *Bernieria*, previously placed in the bulbuls (Pycnonotidae), are yellow and olive in color; the latter with disproportionately long bills. *Thamnornis*, *Randia*, and *Cryptosylvicola* are small with muted coloration, similar to Old World warblers (Sylviidae), in which they were previously placed. *Crossleyia*, *Oxylabes*, and *Hartertula* (formerly *Neomixis*), were considered babblers (Timaliidae) due to their plumage texture and somewhat plumper build. Another species, *Mystacornis crossleyi*, was also traditionally thought to be a Malagasy representative of the babblers, but was recently shown to be part of the endemic Vangidae radiation (Johansson et al. 2008a; Jønsson et al. 2012; Reddy et al. 2012). Confusion in taxonomy has undoubtedly limited our understanding of the ecology and evolution of these and other species on Madagascar.

The tetrakas are part of the sylvioid radiation of the Passeriformes (Johansson et al. 2008), which has undergone several taxonomic rearrangements in the past two decades (Alström et al. 2013; Barker et al. 2004). However, the closest relatives of the Malagasy family remain ambiguous. Most phylogenies place Bernieridae close to the Locustellidae, a recently recognized group of grassbirds, and Donacobiidae, comprised of a single wetland species restricted to South America (Alström et al. 2014; Moyle et al. 2016). Both families are ecologically divergent from the Malagasy group, and suggest a potential shift in the Malagasy lineage from grass/scrub and marsh ancestral habitats to exclusive forest ones. All tetrakas are restricted to thick understory or mid-story conditions, with one species occurring in the canopy. Furthermore, the connection to *Donacobius* is biogeographically anomalous.

In this study, we aim to produce the first comprehensive phylogeny of the Bernieridae (tetrakas) to examine their diversification history on Madagascar, and their place within the avian tree of life. A complete phylogenetic tree is key to helping us understand the dynamics of diversification, as well as the biogeographic origins of endemic Malagasy lineages. To examine the potential for cryptic species within this cryptic family, we sampled all species of Bernieridae from across each of their geographic ranges. We employed integrative taxonomic methods combining phylogenomics of ultraconserved elements (UCEs), genetic clustering of thousands of single nucleotide polymorphisms (SNPs), and analysis of morphometric variation to delineate discrete species. We then reconstructed a robust phylogeny of the Bernieridae and their close relatives using next-generation sequencing of UCEs, producing ∼4500 loci to investigate their evolutionary history.

## Materials and Methods

### Taxon sampling

Our study design included representatives from the Bernieridae, Locustellidae, Donacobiidae and Acrocephalidae families, totaling 130 genotyped individuals. Locustellidae and Donacobiidae have been proposed to be the most closely related families to Bernieridae; however, the evolutionary relationships among the three families are unclear (Alström et al. 2014; Moyle et al. 2016). We therefore aimed to include good representation of all three families in order to better resolve these relationships. Donacobiidae is a monotypic family, containing only *Donacobius atricapilla* (Clements et al. 2018) and we included two subspecies in our analyses. The family Locustellidae has recently been considered to contain 62 species (Clements et al. 2018), however, this figure is probably preliminary e.g. Alström et al. (2018). We included 33 Locustellidae individuals herein, representing 27 species. Three representatives of Acrocephalidae were also included as outgroups, along with one Vangidae as a distant outgroup to root the phylogenies. The full list of non-Bernieridae taxa, including accession numbers, is available in Table S1.

Within Bernieridae, we aimed to assess the presence of cryptic diversity at the species level by sampling each of the 11 recognized species from across their respective geographical ranges. In total, 91 individual Bernieridae have been included. Table S2 presents details on these samples and the supplementary information also includes maps of the sampling locations. Tissue samples used for genotyping are associated with vouchered specimens held at Field Museum of Natural History (FMNH; Chicago), American Museum of Natural History (AMNHDOT; New York), University of Kansas Biodiversity Research Institute (KUBI; Lawrence), Burke Museum of Natural History and Culture (BMNHC; Seattle), Louisiana State University Museum of Natural Science (LSUMZB; Baton Rouge), and Mention Zoologie Biologie Animale, Université d’Antananarivo (UADBA, formerly Département de Biologie Animale, MJR; Antananarivo).

### UCE sequencing

We extracted DNA using QIAGEN DNeasy Blood and Tissue Kits following the manufacturer’s protocol. We prepared UCE libraries for 130 taxa following described methods (Faircloth et al. 2012; McCormack et al. 2013) with minor modifications. Briefly, DNA was normalized to 10 ng/µL and fragmented via sonication (Covaris, Model #M220) to approximately 550 base pairs (bp). Samples were end-repaired and A-tailed, then Illumina TruSeqHT adapters were ligated with either a KAPA Hyper Prep Kit (Kapa Biosystems) or a TruSeq DNA HT Sample Prep Kit (Illumina), following manufacturers’ instructions. Libraries were amplified by limited-cycle (16—18) PCR using Kapa HiFi DNA polymerase (Kapa Biosystems), normalized, and then pooled into sets consisting of eight libraries each with a total of 500 ng of sample. We enriched the pooled libraries for 5060 UCE loci using MYbaits capture kits (Tetrapods 5K v1, Arbor Biosciences) following the manufacturer’s instructions. Enriched libraries were quantified using qPCR (Kapa Library Quantification Kit) and a Qubit Flourometer (Invitrogen), then normalized. Pair-end sequencing was carried out with either 2 x 250 bp or 2 x 150 bp on the Illumina HiSeq2500 platform, or 2 x 300 bp on the Illumina MiSeq v3 platform. DNA sequence reads are archived on NCBI SRA (XXXXX).

### Mitochondrial DNA sequencing

We sequenced mtDNA genes NADH dehydrogenase 3 (ND3) and cytochrome b (CYTB) to facilitate divergence time estimation, because divergence rates in birds are available for these loci (Weir and Schluter 2008) whereas evolutionary rates for UCEs are unknown. We amplified and sequenced ND3 using standard PCR and Sanger sequencing methods with primer pairs ND3-L10751 and ND3-H11151 or ND3-L10755 and ND3-H11151, and CYTB with primers CYTB-L14990 and CYTB-H15916. We used Geneious 9.0.5 for alignment and sequences were deposited in GenBank (TBA — TBA).

### Preparation of UCE alignments

We prepared alignments of UCE loci for phylogenetic analyses using the PHYLUCE 1.5 package (Faircloth 2015). Firstly, the demultiplexed reads were trimmed of adapters and low-quality bases using Illumiprocessor (Faircloth 2013). Trinity 2.0.4 (Grabherr et al. 2011) was then used to assemble contigs, and UCE loci were extracted from among the contigs using PHYLUCE and aligned with MAFFT 7 (Katoh et al. 2002; Katoh and Standley 2013). The alignments were trimmed using the edge-trimming algorithm available in PHYLUCE. Data matrices of either 75% or 90% completeness were generated, where ‘completeness’ refers to the minimum number of taxa sequenced for a locus to be included in the matrix. The specifics of each data matrix are given in the relevant sections below. During the bioinformatics process we also extracted CYTB and ND3 sequences from the off-target contigs using the Megablast function within Geneious 9.0.5, in order to complete our mtDNA datasets.

### SNP calling methods

We prepared datasets of single nucleotide polymorphisms (SNPs) for several species to aid our assessment of cryptic species (see next section). We called SNPs following the protocol of the seqcap_pop pipeline (Harvey et al. 2016), with some modifications. In brief, we used Trinity 2.0.4 to assemble the clean reads across all specimens in a dataset into contigs *de novo*. We extracted contigs matching UCE probes using PHYLUCE and designated these as a reference for SNP calling. The reads for each individual were then mapped to the reference using BWA (Li and Durbin 2009), with a maximum of four mismatches allowed per read. We used Picard (http://broadinstitute.github.io/picard/) and SAMtools (Li et al. 2009) to convert sam files to bam format, soft-clip reads beyond the reference, add read groups for each sample, and merge bam files across all samples in the dataset. We then used the Genome Analysis Toolkit (GATK; McKenna et al. 2010) to realign reads and indels, call SNPs, annotate SNPs and indels, mask indels, and remove SNPs with a quality score < Q30. We then conducted additional filtering of the SNPs using VCFtools 0.1.15 (Danecek et al. 2011): we specified a minimum read depth of three for a genotype call; removed any SNPs with a minor allele count < 2; restricted to biallelic SNPs; and removed any variants not genotyped in 100% of individuals. Finally, we selected one SNP at random per contig to reduce linkage in the final dataset using a custom python script, and then used VCFtools to calculate mean sequencing coverage of each SNP. PGDSpider 2.1.0.0 (Lischer and Excoffier 2012) was used to convert vcf files into other formats required for analysis.

### Detection of cryptic speciation within Bernieridae

We examined each of the 11 species within the Bernieridae family for evidence of cryptic species-level diversity, in order to identify all species in the family before reconstructing its evolutionary history. There have been several recent discoveries of unrecognized species in Malagasy birds (Younger et al. 2019; Younger et al. 2018), and it has been suggested that this may also be the case for some taxa within Bernieridae (Block et al. 2015). In total we sequenced 91 Bernieridae individuals for UCEs. For each species we included representative individuals from all key biogeographic regions inhabited, as far as possible given the availability of tissues. The full methods and results for these analyses are included in the supplementary information, with a breakdown by species (Fig. S1-18).

We first generated a 75% complete data matrix for the full dataset of 130 taxa, including the 91 Bernieridae individuals (4317 UCE loci, total length 3,912,738 bp). We used RAxML 8.2.7 (Stamatakis 2014) to infer a maximum likelihood (ML) phylogeny for this concatenated UCE dataset. We performed rapid bootstrapping analysis and a search for the best-scoring ML tree in a single program run, using the MRE-based bootstopping criterion (Pattengale et al. 2010) to ascertain when sufficient bootstrap replicates had been generated. The search was conducted under the GTR GAMMA site-rate substitution model.

For each species, we asked whether geographically distinct lineages with >90% bootstrap support were evident in the UCE phylogeny (Fig. S1). If so, we then assessed whether those lineages were also supported as distinct with minimal admixture in genetic clustering analyses using SNP data. Distinct phylogenetic lineages were evident for *Bernieria madagascariensis* (n = 20), *Oxylabes madagascariensis* (n = 10), and *Xanthomixis zosterops* (n = 20) (see SI for detailed results), therefore for these three taxa we prepared datasets of single nucleotide polymorphisms (SNPs) as described above and conducted genetic clustering analyses. Based on these combined methods, we found clear genetic evidence of species-level divergences in only one Bernieridae species, *B. madagascariensis*. For *B. madagascariensis* we initially examined 20 individuals and recovered three distinct phylogenetic lineages with 100% bootstrap support (Fig. S2), the same groupings based on genetic clustering analyses (Structure; Pritchard et al. 2000), and no evidence of admixture except for one potentially hybrid individual (Fig. S3). To further examine the possibility that these three lineages represent distinct species, we increased our sample to 39 individuals to allow better delimitation of geographic boundaries of these clades (see below).

### Species delimitation of *Bernieria*

We used a dataset of 39 individuals collected from across the range of *Bernieria* (Fig. 1a) to test the hypothesis that cryptic species are present within the genus. We synthesized data from UCE loci, mitochondrial DNA, and morphometrics in an integrative systematics approach to assess species limits.

**Figure 1.**
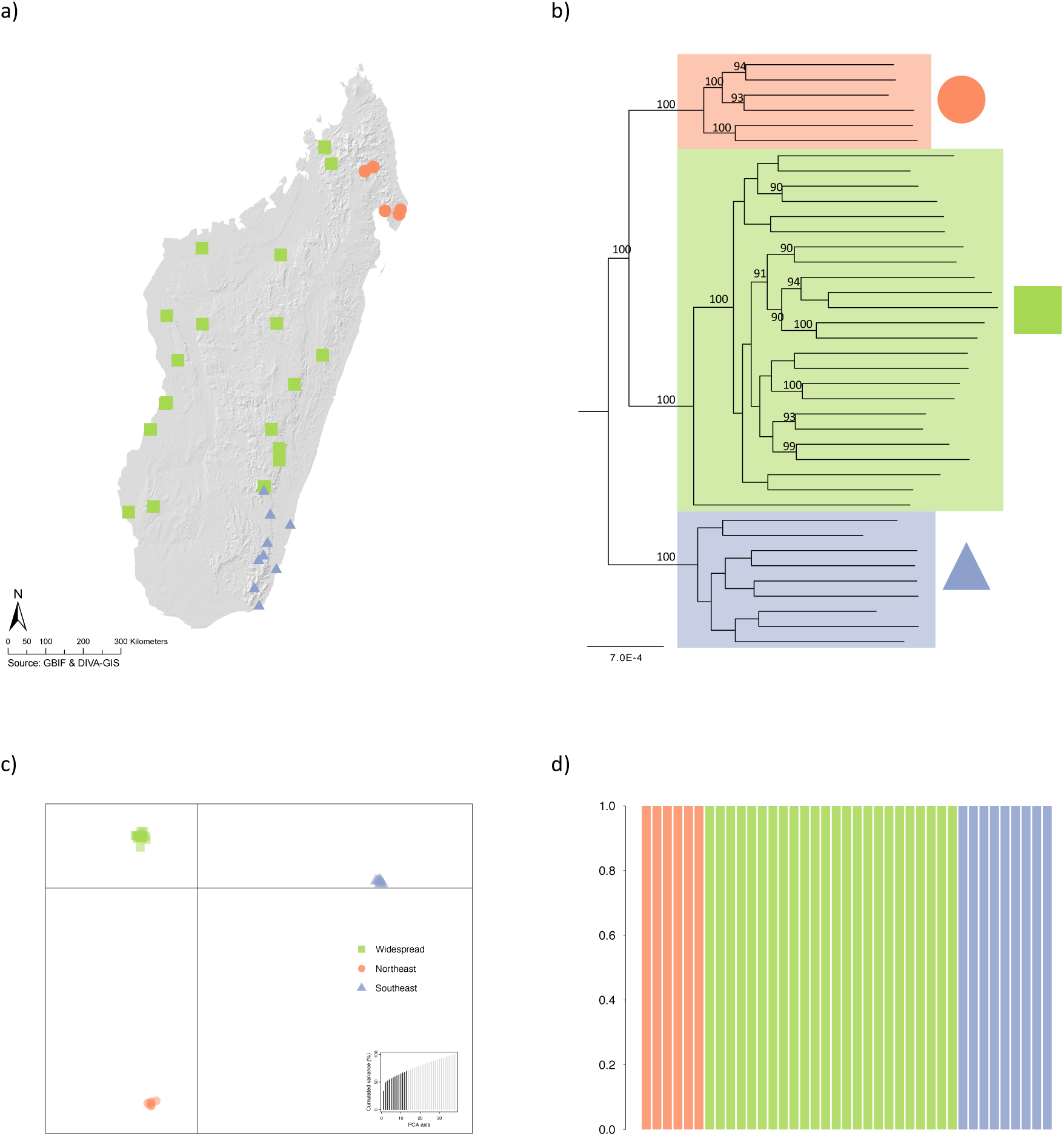
Species delimitation within *Bernieria*. (a) Map of study sampling sites of *Bernieria.* Members of the Widespread clade (WS; n = 24) are indicated by green squares, the Southeastern clade (SE; n = 9) by blue triangles, and the Northeastern clade (NE; n = 6) by orange circles. See Table S1 for latitude/longitude and accession numbers. (b) Phylogenetic relationships within *Bernieria*. Maximum-likelihood phylogeny of 4140 concatenated UCE loci (3,822,981 bp). Support values are shown for nodes that received >70% bootstrap support. (c) Discriminant analysis of principal components (DAPC) for *Bernieria*, based on 3768 unlinked SNPs. Discriminant functions 1 vs. 2, with the amount of variance explained by each principal component (PC) displayed on the bar graphs and the number of PCs retained indicated in black, and (d) the posterior membership probabilities of individuals to each of the three clusters.

### Phylogenetics

We prepared a 75% complete UCE matrix of 39 *Bernieria* individuals plus one outgroup (4140 UCE loci with a total length of 3,822,981 base pairs) and inferred a ML phylogeny using RAxML as described above for the 130 taxa dataset.

### Genetic clustering analyses and summary statistics

We called SNPs for the 39 *Bernieria* individuals (3768 SNPs, 79X mean coverage) and used three different clustering methods to estimate genetic groups: Discriminant Analysis of Principal Components (DAPC) (Jombart, Devillard, & Balloux, 2010), the Bayesian clustering algorithm employed within Structure 2.3.4 (Pritchard, Stephens, & Donnelly, 2000), and principal component analysis (PCA). We implemented DAPC in *adegenet* (Jombart 2008; Jombart and Ahmed 2011). First, we used successive *K*-means clustering to estimate the most likely number of clusters in the dataset, and the assignment of individuals to those clusters. The DAPC method then creates discriminant functions to maximize the variance among, whilst minimizing the variance within, those genetic clusters. Cross-validation with 1,000 replicates was used to determine the optimal number of PCs to retain, testing between 1 and 13 (i.e. up to a third of n = 39). Finally, we plotted the posterior membership probability of all *Bernieria* individuals to the genetic clusters.

Structure (Pritchard et al. 2000) uses a Bayesian clustering method with a Markov Chain Monte Carlo (MCMC) sampling procedure to estimate membership coefficients for each individual to each of a given number of clusters (*K*). We performed Structure analyses for the *Bernieria* SNP dataset using the admixture model with correlated allele frequencies and without sampling location priors. We carried out an initial run of 100,000 generations, discarding the first 50,000 as burn-in, with *K* = 1 and lambda allowed to vary in order to estimate a value for lambda (the allele frequencies prior). We then set the value of lambda to the estimated value for subsequent runs. For the main analyses, we varied the number of clusters from *K* = 1 to *K* = 6, and ran each analysis for 700,000 generations with the first 200,000 discarded burn-in. The analysis for each value of *K* was repeated ten times. We used Structure Harvester Web 0.6.94 (Earl 2012) to assess convergence across replicates, to determine the most optimal value(s) of *K* for the dataset based on the log likelihood of each value of *K* and the Evanno method (Evanno et al. 2005), and to prepare input files for CLUMPP 1.1.2 (Jakobsson and Rosenberg 2007). We used CLUMPP to average membership coefficients across the replicates. Finally, Distruct 1.1 (Rosenberg 2004) was used to visualize the results for *K* = 2, *K* = 3, and *K* = 4. We chose to visualize three values of *K* in order to gain insight into potential levels of genetic structure within *Bernieria*.

We performed PCA for *Bernieria* using the SNP dataset. We used the *scaleGen* function in *adegenet* to scale and centre allele frequencies, and then performed PCA using the *dudi.pca* function from the *ade4* v1.7-11 package. We plotted the first two PCs only, as these clearly differentiated the three genetic groups.

We used Genodive 2.0b27 (Meirmans & Van Tienderen, 2004) to calculate the Weir and Cockerham unbiased weighted *F_ST_* estimator (Weir & Cockerham, 1984) between each pair of clades, with significance calculated using 10,000 permutations of the data. This measure has been shown to be robust to small sample sizes, even when *F_ST_* is low (Willing, Dreyer, & van Oosterhout, 2012).

### Divergence time estimation

We estimated divergence times among *Bernieria* lineages using time-calibrated Bayesian phylogenetic analyses within BEAST 2.4.4 (Bouckaert et al. 2014). We used mtDNA sequences (ND3 and CYTB) because estimates of divergence rates in birds are available for these loci (Weir and Schluter 2008), whereas evolutionary rates for UCEs are unknown. The mtDNA gene trees resolved the same three lineages as the UCE dataset, with 100% support for each. We partitioned the data into ND3 and CYTB and specified the nucleotide substitution models as HKY for both genes to reflect the optimal models as estimated with PartitionFinder 2 (Lanfear et al., 2016). We used the Yule tree prior with a strict molecular clock. We calibrated the analysis using the divergence rate of CYTB for Passeriformes of 2.07% (± 0.20) per million years (Weir and Schluter 2008) as a reference rate (lognormal, mean = 0.01035, SD = 0.05). We ran two independent analyses for 120 million generations each, and then confirmed convergence of the posteriors using Tracer v1.6 (Rambaut and Drummond 2007). We estimated a maximum clade credibility tree with mean node heights from the posterior after removing the first 10% of samples as burn-in.

### Morphological variation

To determine whether genetic lineages of *Bernieria* were morphologically differentiated, one of us (NLB) measured 52 adult skin specimens (26 each of males and females; Table S5) housed in the FMNH for six standard avian characters: bill length from the crown to tip (BL), bill width at nares (BW), bill depth at nares (BD), tarsus length (TL), tail length (Tail), and wing length (WL). These measurements followed descriptions in (Baldwin et al. 1931). We used Mitutoyo Digital Calipers to measure bill and tarsus to an accuracy of 0.01 mm. For wing and tail lengths, we used a wing rule to an accuracy of 1 mm. We repeated all measurements three times, checked for outliers (by confirming that all measurements for an individual were within one standard deviation), and then averaged values for subsequent analyses. The summary of these measurements is in Table S5.

For multivariate analyses, we scaled and log-transformed the measurement data; as two specimens were missing data, we removed these from the multivariate dataset. First, we tested for sexual dimorphism by using ANOVAs to compare the measurements of males and females of each genetically identified clade. Since most measurements showed significant sexual dimorphism, we conducted all subsequent analyses separately for each sex. Next, we performed a principal components analysis (PCA) to examine morphological variation among *Bernieria* and to determine if the genetically identified groups represent distinct clusters in morphospace. We used MANOVA to determine if the centroids of each group were statistically different.

### Family-level evolutionary relationships

We used a pruned dataset of 51 taxa, wherein each Bernieridae species was represented by a single individual, to reconstruct evolutionary relationships within and among Bernieridae, Locustellidae, Donacobiidae, and Acrocephalidae. We prepared both 75% and 90% complete data matrices of these taxa. We inferred ML phylogenies using RAxML 8.2.7 (Stamatakis 2014). We performed both partitioned and unpartitioned concatenated analyses for the 75% complete data matrix, and an unpartitioned concatenated analysis for the 90% complete data matrix. To estimate an appropriate partitioning scheme we used the Sliding-Window Site Characteristics (SWSC) entropy based method (Tagliacollo and Lanfear 2018) to generate subsets of each locus based on within-locus heterogeneity (e.g., the UCE flanking regions are typically more variable than the conserved core). These subsets were then passed to PartitionFinder 2 (Lanfear et al. 2014; Lanfear et al. 2016) to estimate a partitioning scheme for phylogenetic analysis by grouping together similar subsets. For the RAxML analyses, we performed rapid bootstrapping analysis and a search for the best-scoring ML tree in a single program run, using the MRE-based bootstopping criterion (Pattengale et al. 2010) to ascertain when sufficient bootstrap replicates had been generated. The searches were conducted under the GTR GAMMA site-rate substitution model.

We also inferred a phylogeny under the multispecies coalescent method. For this analysis we used a subset of the 25% of UCE loci with the greatest number of parsimony informative sites, consisting of 1070 UCE loci with 122–331 parsimony informative sites each. We estimated each gene tree with 100 ML searches under GTR GAMMA using RAxML, and then reconciled these 1070 phylogenies into a gene tree-species tree using ASTRAL 4.10.12 under the default settings (Mirarab and Warnow 2015).

The evolutionary relationships among Locustellidae, Bernieridae, and Donacobiidae have a history of phylogenetic conflict. We inferred phylogenies using a single representative per family to see if reduced taxon sampling would alter our estimates of these relationships. We prepared 75% and 100% complete data matrices for a 4-taxon dataset of *Locustella fluviatilis* (Locustellidae), *Oxylabes madagascariensis* (Bernieridae), *Donacobius atricapilla atricapilla* (Donacobiidae), and *Acrocephalus orientalis* (Acrocephalidae). These matrices contained 4299 and 2826 UCEs, respectively. We inferred concatenated ML phylogenies for these using RAxML, as described previously.

### Lineage diversification rates within Bernieridae

We estimated speciation rates over time within the Bernieridae, to determine whether the diversification mode was consistent with a model of adaptive radiation driven by ecological opportunity (Losos and Mahler 2010; Schluter 2000). We first performed a time-calibrated Bayesian phylogenetic analyses on mtDNA sequences (ND3 and CYTB) using BEAST 2.4.6 (Bouckaert et al. 2014). We used a dataset of 49 taxa, including all 13 species of Bernieridae and representatives from each of the other three families. mtDNA genes were used because estimates of divergence rates in birds are available for these loci (Weir and Schluter 2008). The data was partitioned into CYTB and ND3, with substitution models specified as GTR with four gamma categories, and TN93 with four gamma categories, respectively, to reflect the optimal models selected by PartitionFinder 2 (Lanfear et al., 2016). We used a lognormal relaxed clock model, calibrated with the divergence rate of CYTB for Passeriformes of 2.07% (± 0.20) per million years (Weir and Schluter 2008) as a reference rate (lognormal, mean = 0.01035, SD = 0.07). We also calibrated the MRCA of Bernieridae, Locustellidae, Donacobiidae and Acrocephalidae as 17.42 Ma, with a 95% confidence interval of 16.13 – 18.97 Ma (normal distribution, mean = 17.42, sigma = 0.87) in accordance with a recent dating study that included these groups (Moyle et al. 2016); note that there are no relevant fossils attributed to these families to aid our estimates. The topology was constrained to match our UCE phylogeny, and two independent analyses were performed to ensure reproducibility of the posteriors. The MCMCs were run until convergence, confirmed by inspection of the posterior distributions and effective sample sizes in Tracer v1.6 (Rambaut and Drummond 2007). We estimated a maximum clade credibility tree with median node heights from the posterior after removing the first 10% of samples as burn-in.

Prior to diversification rate analyses we removed the outgroup Acrocephalidae taxa and pruned the time-calibrated phylogeny to species-level sampling (i.e. additional subspecies were removed). This time-calibrated phylogeny was then used to calculate the Pybus and Harvey gamma (γ) statistic (Pybus and Harvey 2000) for the Bernieridae clade, using the ape package (Paradis and Schliep 2018) in R. γ provides a measure of the distribution of speciation times in a phylogenetic tree relative to our expectation under a pure-birth model of constant-rate speciation. γ = 0 indicates a constant rate of speciation over time, whereas γ < 0 and γ > 0 indicate declining and increasing rates of speciation over time, respectively. To determine the effect (if any) of cryptic species on our estimate of speciation rate constancy over time, we removed the two new *Bernieria* species and recalculated the statistic. We also estimated speciation rates of the Bernieridae clade over time using BAMM 2.5 (Rabosky 2014). We specified the fraction of sampling per clade, set the prior on number of rate shifts to 1, and ran the analysis for 30 million generations. We assessed convergence and plotted the speciation rate over time for the Bernieridae clade using Bammtools 2.1 (Rabosky et al. 2014). Note that we did not attempt to detect rate shifts given the relatively small size of our dataset (n = 42 tips).

## Results

### Genetic data

Our UCE capture protocol recovered an average of 4180 UCE loci per individual, with 4982 UCE loci recovered across all 130 taxa. We prepared several datasets from these UCE loci. The 75% complete matrix for the full 130 taxa contained 4317 loci and was 3,912,738 bp in length. For the 51 taxa dataset, we selected one representative per species with the greatest amount of UCE sequence recovered. For this taxon set we prepared both 75% and 90% complete matrices, consisting of 4281 loci (4,073,553 bp) and 2489 loci (2,423,537 bp), respectively. The *Bernieria* 75% complete matrix contained 4140 UCE loci (3,822,981 bp). Our SNP calling and filtering protocols generated high-coverage, unlinked SNP datasets for *Bernieria* (n = 20), *Bernieria* (n = 39), *O. madagascariensis* (n = 10), and *X. zosterops* (n = 20). Mean sequencing depth per SNP ranged from 62 to 119, with the number of SNPs per dataset ranging from 2668 to 3768.

### Integrative species delimitation within Bernieridae

We evaluated species limits across Bernieridae and found convincing evidence of cryptic species-level diversity in *Bernieria* only. Both *Oxylabes madagascariensis* and *Xanthomixis zosterops* have apparent genetic population structure in the eastern humid forest. In the case of *O. madagascariensis*, there is some evidence for two genetic populations in the north and south respectively, but our results suggest admixture between these, and our clustering analyses gave almost equal support for either one or two genetic clusters (Fig. S8-11). Our results for *Xanthomixis zosterops* suggest the presence of three genetic populations, but many individuals have evidence of mixed ancestry and our clustering and phylogenetic analyses did not assign individuals to the same lineages (Fig. S16-18), therefore these divergences are most likely population rather than species level differences. We suggest that both of these species should be further investigated with geographically comprehensive sampling in the future to delimit the boundaries of populations. Interestingly, other taxa with similar distributions throughout the eastern humid forests showed no evidence of genetically distinct lineages in our analyses (e.g. *Hartertula flavoviridis*, *Xanthomixis cinereiceps*). A thorough description of the population genetic and species delimitation analyses for all tetraka taxa other than *Bernieria* are presented in the Supplemental Information (Fig. S1-18).

Our ML phylogeny of *Bernieria* resolved three clades with 100% bootstrap support each (Fig. 1b). These clades were correlated with geography; one was found in the northeast (NE), one in the southeast (SE), and the other was widespread in western Madagascar and the central portions of the eastern forest (WS; Fig. 1a). Our BEAST analysis of mtDNA sequences had posterior supports of 1.0 for each of the same three clades, however, one individual (FMNH 427373 collected just south of Ranomafana National Park and the voucher specimen saved as a skeleton) was in the SE mitochondrial clade, but the WS UCE clade, indicating that it is likely a hybrid. The relationships among the three mitochondrial clades differed from the UCE phylogeny; the NE and SE mitochondrial lineages appear as sister taxa, whereas the NE and WS UCE clades are sisters. Mito-nuclear biogeographic discordance such as this could be the result of historical secondary contact between two of the clades (Toews and Brelsford 2012), a hypothesis that could be examined in future studies. Using mitochondrial genes, we estimated that the NE and SE mitochondrial clades diverged approximately 3.10 Ma (median estimate, 95% HPD: 2.50 – 3.82 Ma), with the split from the WS clade occurring around 4.84 Ma (median estimate, 95% HPD: 4.04 – 5.72 Ma).

We used the dataset of 3768 unlinked SNPs (mean coverage 79X) to perform genetic clustering analyses of *Bernieria*. Successive *K*-means clustering clearly indicated *K* = 3 as the optimal number of clusters based on the Bayesian Inference Criterion (Fig. S19), and DAPC was able to differentiate among these with 100% support (root mean squared error = 0) (Fig. 1c). All individuals were assigned to their respective clusters with a posterior membership probability of 1.0, indicating no evidence of admixture (Fig. 1d). In our Structure analysis, the maximum posterior log probability of the data was achieved at *K* = 3 (Fig. S20), but the rate of change in log probability (deltaK, Evanno method) was greatest at *K* = 2. We therefore plotted membership probabilities for *K* = 2 – 4 (Fig. S21). Assignments of individuals to these clusters were consistent with the results of our phylogenetic analyses. In the three-cluster scenario, all individuals in the NE and SE clades were assigned with 100% support. In the WS clade, FMNH 427373 appears to be a hybrid of 0.73 WS and 0.27 SE ancestry, consistent with its divergent placement in the mitochondrial and nuclear phylogenies. In the two-cluster scenario, the WS cluster appears most distinct, with NE and SE individuals generally clustering together (Fig. S21). No additional genetic structure was revealed by *K* = 4 (Fig. S21). For our PCA, the first two PCs explained 13.14% and 9.65% of the variation in the data, respectively. The three genetic lineages were well separated on a plot of PC1 vs. PC2, with no overlap, and the mixed ancestry of the hybrid individual is apparent (Fig. S22). Our estimates of *F_ST_* between pairs of clades ranged from 0.422 to 0.454 (*p*-values < 0.0001), suggesting strong, statistically significant genetic differentiation among the three clades (Table S3-4).

The three genetically distinct clades are differentiated by subtle differences in morphological variation. The PCAs of each sex resulted in six principal components with the first four encompassing more than 90% of the variance in males (Fig. S23) and 95% in females (Fig. S24). Although the plots of PC variance showed some overlap of the three genetic lineages in 2-D morphospace, the MANOVA indicated there is statistically significant morphological differences among them in multidimensional space (Fig. S23-24). Pairwise ANOVA (t-tests) for the six morphological characters also showed significant differences between all pairs of genetic groups (Table S6). The SE and WS genetic groups were the most morphologically divergent from each other, with significant differences in tarsus length, tail length, and bill width of females and wing and bill width of males. In comparison to those from the SE group, the WS birds were smaller, but with longer tails. Between the NE and SE genetic groups, there were two statistically significant differences – tarsus length of females and wing length of males. There was only one significant difference between NE and WS, in the tail length of males.

Geographically, based on our current sampling, the NE *Bernieria* clade is allopatric, and separated from the WS clade by the northern Tsaratanana Massif (Fig. 1a). The WS and SE clades are geographically separated by just seven kilometers in the eastern humid forest, at the Ambositra-Vondrozo Forest Corridor Reserve and Parc National Andringitra, respectively. However, the clades have minimal overlap in elevational distribution in this region. Based on current data, the SE clade is found between 50 and 1055 m (mean = 527 m), and the WS clade in the eastern forest is found between 980 and 1710 m (mean = 1300 m), suggesting a narrow elevational contact zone of < 100 m. Interestingly, the WS clade is found at elevations from 18 to 1517 m in the western part of its range, suggesting that its restriction to montane habitats in the eastern humid forest may be a result of competition with the SE clade, rather than adaptation to montane forest or physiological limitations.

Overall, our analyses provide strong evidence for three genetically divergent and phenotypically distinct lineages of *Bernieria* that originated during the Pliocene. There is one apparent case of hybridization between two of these lineages, but no other admixture. The lineages warrant recognition as distinct species, and as such we have included all three species in our family-level phylogenetic and diversification rate analyses. The current taxonomy of *B. madagascariensis* includes two subspecies; *B. m. inceleber* (Bangs and Peters 1926), inhabiting western dry forest, spiny bush, and the humid forest of Mt d’Ambre in the far north, and *B. m. madagascariensis* (Gmelin 1789), occupying other humid forest formations. *B. madagascariensis* was originally described as *Musciapa madagascariensis* (Gmelin 1789) based on specimens collected at Tolagnaro (Stresemann 1952), within the range of the SE clade. We therefore propose that this clade should retain the *B. madagascariensis* species name, and that the name *B. inceleber* should be applied to the WS clade. We suggest the common name ‘Obscure Bernieria’ for *B. inceleber*, to reflect its Latin name. The third species, corresponding to the NE clade, referred to here as *Bernieria sp.*, will be formally described and named in a separate manuscript.

### Family-level evolutionary relationships

Our phylogenetic analyses converged on a strongly supported topology for Bernieridae, Locustellidae, Donacobiidae, and Acrocephalidae (Fig. 2). In our ML analyses, every bipartition had 100% bootstrap support and the topology was entirely consistent across the 75% and 90% complete data matrices, and partitioned and unpartitioned analyses (Fig. S25-26). The ASTRAL species tree resolved the same topology and had a normalized quartet score of 0.93 (Fig. S27). Almost every quadripartition had support of 1.0, except for two of the shallow nodes within *Locustella*.

**Figure 2.**
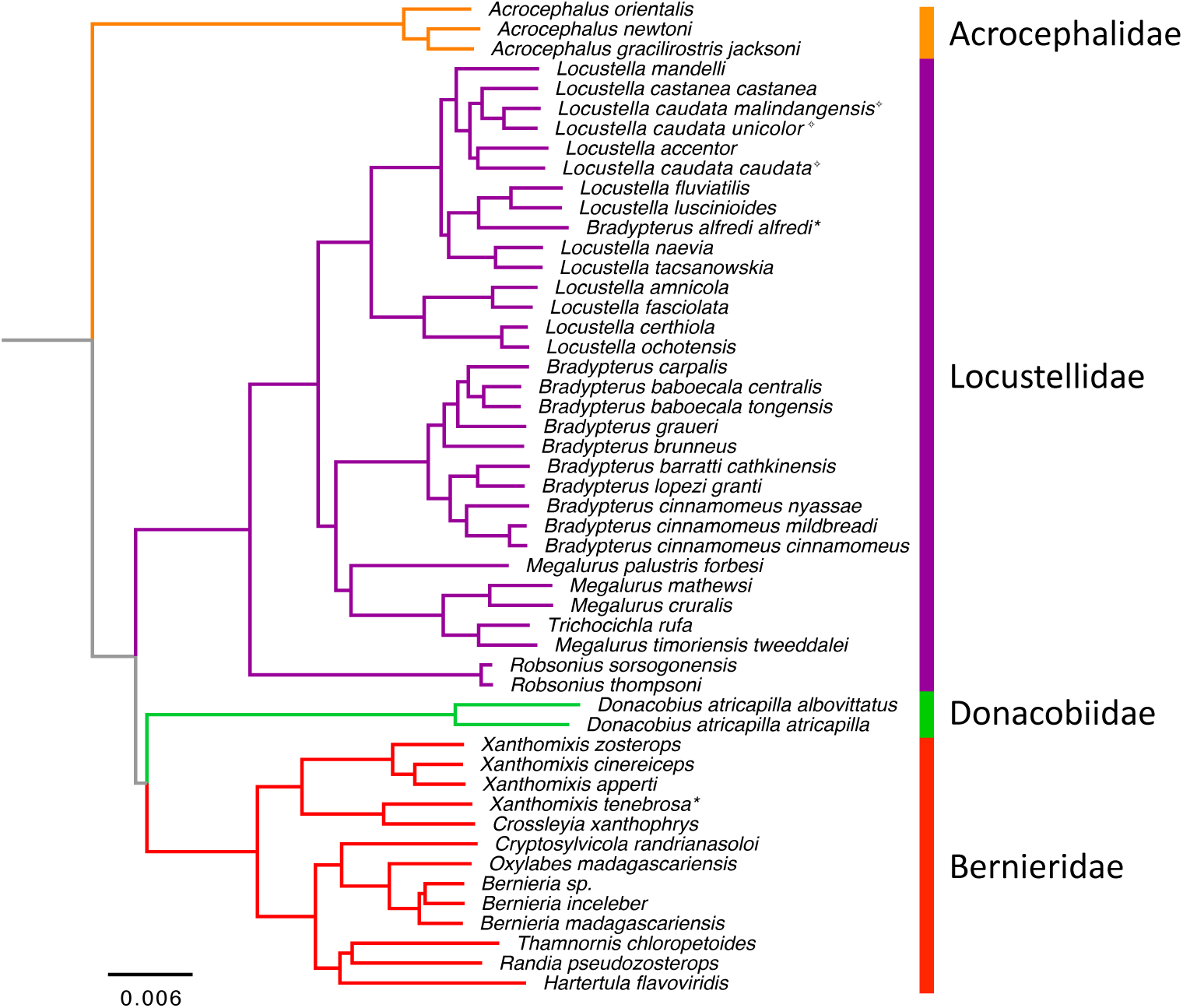
Evolutionary relationships among Bernieridae, Locustellidae, Donacobiidae, and Acrocephalidae. Partitioned maximum-likelihood phylogeny of the 75% complete data matrix containing 4281 concatenated UCE loci (4,073,553 bp). All branches have 100% bootstrap support. * indicates species that should be reclassified at the generic level; ✧ indicates paraphyletic species.

All our analyses show that the Bernieridae are a monophyletic radiation endemic to Madagascar (Fig. 2,4). This includes the ten species in the original designation of the family (Cibois et al. 2010), as well as *Randia pseudozosterops*, the elevation of the form *Bernieria. m. inceleber* to a full species, and the new *Bernieria sp.*, for a total of 13 species in the family. *Xanthomixis* is paraphyletic with respect to *Crossleyia* in all our analyses. Given the depth of the divergence of *X. zosterops*, *X. cinereiceps*, and *X. apperti* from *X. tenebrosa* and *C. xanthophrys*; *X. tenebrosa* is better regarded as a member of the *Crossleyia* genus.

**Figure 3.**
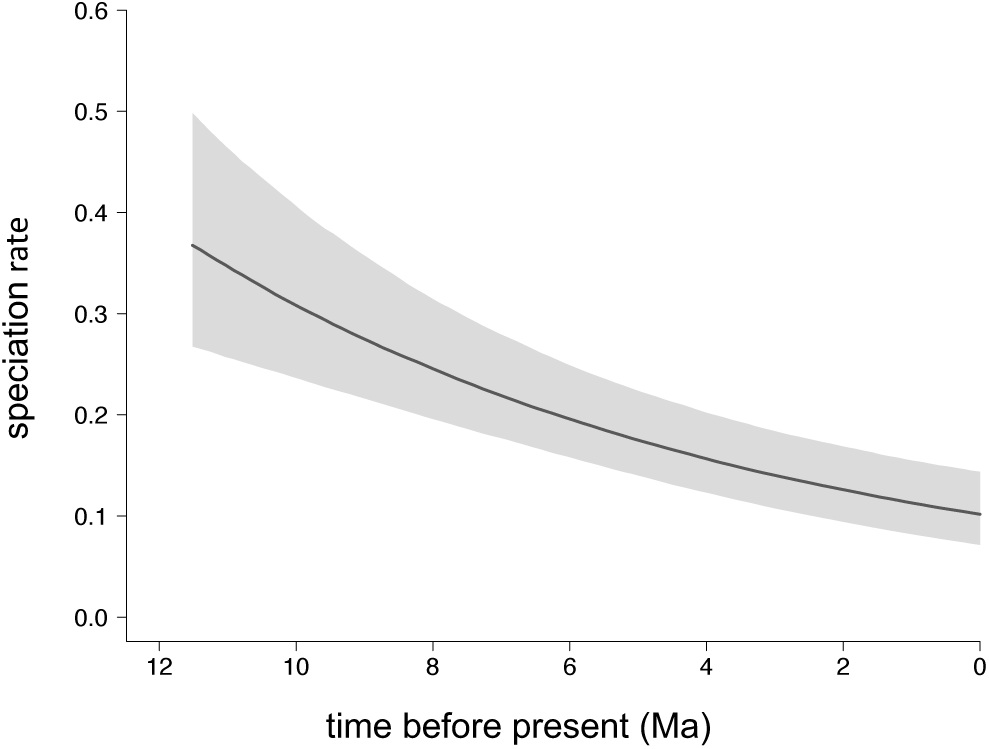
Speciation rate of the Bernieridae clade over time. (per lineage per Ma), where solid line indicates mean estimate and shading represents the 95% highest posterior density interval.

**Figure 4.**
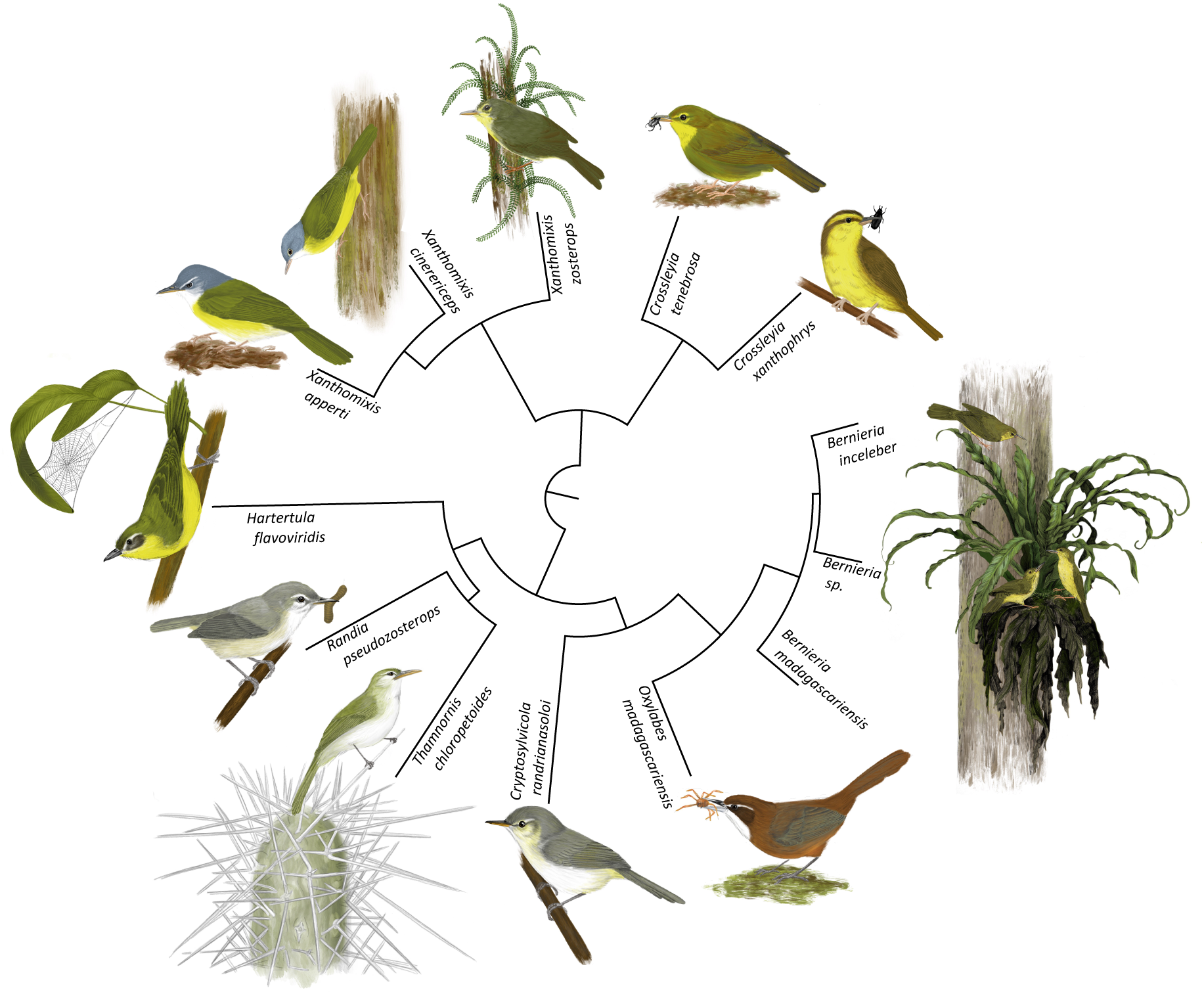
Evolutionary relationships within the Bernieridae. Illustrated to show species in their typical habitats. Partitioned maximum-likelihood phylogeny of the 75% complete data matrix as in Figure 2; all branches have 100% bootstrap support.

Among the four families we found Acrocephalidae to be the most divergent. Donacobiidae was resolved as the sister family to Bernieridae with 100% support, yet the corresponding branch is notably short, suggesting that the divergences of Locustellidae, Bernieridae, and Donacobiidae from their common ancestor may have been rapid. This relationship was the same when only a single taxon per family was used; *Oxylabes* and *Donacobius* were monophyletic with 100% bootstrap support in our ML analyses of the four-taxon dataset.

Our sampling of Locustellidae included 27 of the 62 currently recognized species (Clements et al. 2018). We found four major clades within this sampling. The *Robsonius* genus was most divergent. The other three clades correspond to *Locustella*, *Bradypterus*, and *Megalurus*, and we found several inconsistencies with the existing taxonomy in these groups. *Bradypterus alfredi* falls within our *Locustella* clade, with 100% support in all analyses, and the three subspecies of *Locustella caudata* included here are paraphyletic with respect to *Locustella accentor*, *Locustella castanea*, and *Locustella mandelli*.

### Bernieridae speciation rates

Reconstruction of Bernieridae speciation rates over time shows a clear slowdown in speciation, with the most rapid diversification occurring at the onset of the radiation *ca*. 11– 12 Ma (Fig. 3). The γ statistic for Bernieridae is −3.58, a highly unlikely observation if speciation rates were constant over time (*p* < 0.001) indicating strong support for declining rates of speciation within the family. When we removed *Bernieria inceleber* and *Bernieria* nov. sp., γ was −3.29 and still highly statistically significant (*p* < 0.001), indicating a strong signal of slowdown in speciation rate regardless of whether cryptic species are included in the phylogeny. The decline of speciation rate in the Bernieridae clade over time is consistent with the ecological opportunity model, whereby speciation rates are expected to be high upon the colonization of a new environment (in this case, Madagascar), and then slow over time as niches fill and a new equilibrium is reached (Losos and Mahler 2010; Schluter 2000).

## Discussion

Our study presents the first complete phylogeny of the recently recognized Bernieridae family. Critical to our understanding of the evolution of any family is a characterization of its species composition. Here, we used an integrative taxonomic approach combining phylogenomics of UCEs, genetic clustering of thousands of SNPs, and statistical analyses of morphology, to examine species limits across Bernieridae and facilitate a detailed reconstruction of its evolutionary history. We uncovered two cryptic species within this cryptic family, expanding the number of recognized Bernieridae species to 13. We then utilized a large-scale genomic dataset with comprehensive taxon sampling to reconstruct relationships within the family and its place within the avian tree of life, resolving a long-standing issue of phylogenetic conflict among the Bernieridae, Locustellidae, and Donacobiidae families.

### Bernieridae: a cryptic radiation

The Bernieridae family has a fascinating taxonomic history. The original classifications of these “Malagasy warblers” scattered them across three different families on the basis of plumage and morphology, until a mitochondrial study showed that nine taxa of Malagasy passeriforms were united in a single clade endemic to Madagascar (Cibois et al. 2001). This long-overlooked cryptic radiation was only formally described as the Bernieridae in 2010 (Cibois et al. 2010). Ten species were included in the description, and *Randia pseudozosterops* was flagged as a potential addition pending further study. Ours is the first published phylogeny to include all recognized species, the first to include *Randia*, and by far the largest genetic dataset used to analyze the family (Alström et al. 2013; Cibois et al. 2001; Fregin et al. 2012; Johansson et al. 2008b). Our results conclusively show that Bernieridae is a monophyletic radiation endemic to Madagascar and that *Randia* is a member of this radiation. Our phylogeny is also the first to include all *Xanthomixis* and *Crossleyia* taxa in a single analysis, and therefore show that *X. tenebrosa* is best placed within the genus *Crossleyia*.

The significant decline in the rate of Bernieridae speciation over time is suggestive of progressive niche filling, consistent with a model of adaptive radiation driven by ecological opportunity (Losos and Mahler 2010; Rabosky and Glor 2010; Schluter 2000). While the Bernieridae are all insectivores, they are nevertheless ecologically and morphologically diverse (Fig. 4). Morphologically, the species display diversity in their body size, bill shape, tarsus length, and wing and tail proportions (Schulenberg 2003). Their habitat preferences range from humid forest to sub-arid spiny bush. Within a given forest formation, up to seven sympatrically occurring species show stratification – occupying different portions of the understory to canopy, as well as along elevational gradients (Goodman and Putnam 1996; Schulenberg 2003), and up to six species have been observed in the same mixed-species foraging flocks (Schulenberg 2003). These patterns are suggestive of partitioning of ecological niche space consistent with adaptive radiation, making the Bernieridae the second largest avian adaptive radiation (after the Vangidae) on the island of Madagascar. The island’s complex landscape heterogeneity coupled with its geographic isolation since before the origin of birds (Storey et al. 1995) would have provided ample opportunities for diversification of these two adaptive radiations (de Wit 2003; Wilmé et al. 2006). Our reconstruction of the evolutionary relationships within Bernieridae sets the stage for future studies of ecological and phenotypic diversification across this family.

### Bernieridae in the avian tree of life

Hidden clades, as exemplified by the discovery of the Bernieridae, are key missing puzzle pieces in our understanding of the avian tree of life (Reddy 2014). This rewriting of evolutionary history also has important implications for Southern Hemisphere biogeography. Previous work placed Bernieridae within the superfamily Sylvioidea (Johansson et al. 2008b), and in a monophyletic clade with Locustellidae and Donacobiidae (Alström et al. 2014; Fregin et al. 2012; Johansson et al. 2008b; Moyle et al. 2016; Olsson et al. 2013). However, these studies reported conflicting relationships among the three families. Phylogenies based on a combination of nuclear introns, exons, and mitochondrial genes generally recovered Donacobiidae as the sister family to Bernieridae with between 58% and 95% bootstrap support (Alström et al. 2014; Fregin et al. 2012; Olsson et al. 2013), with the exception of one study that reported conflicting topologies from different inference methods (Johansson et al. 2008b), and another that recovered Locustellidae as sister to Bernieridae with low support (Oliveros et al. 2012). A more recent study based on UCEs found conflicting relationships from different inference methods; in a concatenated ML analysis of a 75% complete data matrix Donacobiidae and Locustellidae were monophyletic with 100% support, whereas an ML analysis of the 100% complete matrix, as well as a multispecies coalescent based analysis (SVDquartets), both recovered Donacobiidae and Bernieridae as monophyletic with strong support (Moyle et al. 2016).

Ours is the most complete study of these relationships to date, both taxonomically and genetically, and all of our ML and multispecies coalescent based analyses found Donacobiidae and Bernieridae to be sister families with 100% support. The conflict between our result and that of the previous UCE study is of interest, because both studies targeted the same 5060 UCEs. A key difference is taxon sampling; Moyle et al. (2016) included only a single representative per family, whereas we included many taxa per family. We therefore selected a single representative per family from our dataset, using the taxa most closely related to those in Moyle et al., but still recovered Donacobiidae and Bernieridae as sister taxa with 100% support in our analysis of both 75% and 100% complete data matrices, suggesting the incongruence may instead be related to missing data in their 75% complete matrix. Overall, the relationships among Bernieridae, Donacobiidae, and Locustellidae are stable across all of our analyses, and are consistent with a majority of the previous studies. It seems likely that the divergences among these three families were rapid given the previous phylogenetic conflict, and as evidenced by the gene tree discordance detected in our multispecies coalescent analysis (Fig. S27).

The sister relationship between Donacobiidae and Bernieridae is intriguing from a biogeographic perspective. Donacobiidae is represented by a single species, comprising four different geographical forms (subspecies) and restricted to Central and South America – half the world away from Madagascar. On the other hand, Locustellidae is a large family found in most major bioregions of the Old World, including Madagascar, Africa, the Palaearctic, Southern Asia, and Australasia. A recent reconstruction of the biogeographic history of oscine passerines suggested that the Miocene ancestor of these three families most likely lived in East and/or South Asia (Moyle et al. 2016). Given the far-flung contemporary distributions of Donacobiidae and Bernieridae, this scenario seems as plausible as any.

### Cryptic speciation

Our analysis of species limits across Bernieridae uncovered two unrecognized cryptic species within the previously monotypic genus, *Bernieria*. We found strong support for three deeply divergent genetic lineages, dating to the Pliocene, that are also phenotypically distinct and therefore merit recognition as separate species. Two of these species, *B. madagascariensis* and *Bernieria* sp., have restricted geographical ranges indicative of fine-scale endemism, adding to the growing body of evidence that microendemism in birds may as yet be underestimated on Madagascar (Younger et al. 2018).

With this comprehensive understanding of species limits we were able to reconstruct the evolutionary relationships among all species in the Bernieridae and show that the slowing of speciation rate over time is consistent with adaptive radiation, rather than an artefact of missing species. *Bernieria inceleber* and *Bernieria* sp. join other recent discoveries of cryptic species-level diversity in Madagascar’s endemic avifauna (Younger et al. 2019; Younger et al. 2018). When considered in combination with the recent recognition of the Bernieridae (Cibois et al. 2010), the limitations to our understanding of avian biodiversity becomes apparent. Knowledge of areas of endemism is particularly important for setting conservation priorities, and in light of the immediate threat posed to the island’s forest-dwelling fauna by habitat destruction (Vieilledent et al. 2018), there is an urgent need to examine the other avian families on Madagascar for overlooked species.

The humid forests of Madagascar support ten species of Bernieridae, underlining the importance of this habitat type in the evolution of this adaptive radiation. Interestingly, a previous study of *Xanthomixis zosterops* found four deeply divergent mitochondrial lineages in the eastern humid forests (3.5 – 7.6 % cytochrome *b* divergence; Block et al. 2015). Our analyses of nuclear variation for *X. zosterops* from throughout the humid forests were discordant with these mitochondrial lineages and suggested weak population structure rather than species-level divergences (see Supplementary Information). Biogeographic mito-nuclear discordance such as this has many possible drivers (Toews and Brelsford 2012), ranging from incomplete lineage sorting (Zink and Barrowclough 2008) to secondary contact (Carling and Brumfield 2008), or selection on mitochondrial DNA (Irwin 2012). Future phylogeographic analyses could elucidate the cause of mito-nuclear discordance in *X. zosterops* and yield new insights into the role of the eastern humid forests in generating avian diversity.

### Insights into Locustellidae taxonomy

The Locustellidae (grassbirds and allies) currently comprises 62 species in 11 genera (Clements et al. 2018) found across the Old World. The taxonomy of this family is a work in progress; a recently published phylogeny, based on five loci for 59 of the 62 species, suggested numerous changes (Alström et al. 2018). Our sampling of Locustellidae is only 45% complete at the species level (28/62), but our UCE dataset of more than 4 million base pairs is expansive compared to previous sequencing efforts, leading to several new insights into the evolution of this family.

The three subspecies of *Locustella caudata* (*L. c. caudata*, *L. c. malindangensis*, and *L. c. unicolor*) are paraphyletic in our analyses with respect to both *L. accentor* and *L. castanea*. The two subspecies from Mindanao in the southern Philippines, *L. c. malindangensis* and *L. c. unicolor*, formed a monophyletic clade with *L. castanea* from Sulawesi, whereas the other subspecies, *L. c. caudata* from Luzon in the northern Philippines, was sister to *L. accentor* from Borneo. Recognition of the different subspecies of *L. caudata* as full species may be warranted, pending integrative species delimitation. These would join several other recent discoveries of hidden diversity in the endemic birds of the Philippines (Hosner et al. 2013; Hosner et al. 2018).

The relationships within our *Locustella* clade are broadly in agreement with those of Alström et al. (2018); in particular, *Bradypterus alfredi* was placed within *Locustella* with 100% support in all our analyses, and we second Alström’s suggestion for its reclassification. Within the *Bradypterus* genus, our phylogeny conflicts with Alström’s regarding the clade containing *B. carpalis*, *B. baboecala centralis*, *B. baboecala tongensis,* and *B. graueri*; the relationships within this clade were poorly supported in Alström’s phylogeny (< 60% bootstrap support), suggesting that our additional sequencing effort was necessary to resolve these relationships.

## Concluding remarks

The Bernieridae family went unrecognized by science for decades. Our study resolves the evolutionary history of this cryptic radiation and its place within the avian tree of life, representing an important step forward in understanding the origins and diversity of Madagascar’s endemic avifauna. These results highlight the importance of complete taxon sampling for phylogenetic studies, including the identification of cryptic or hidden species. A comprehensive appreciation of species-level diversity is crucial for understanding evolutionary processes, and we advocate that evolutionary biologists consider an integrative species delimitation approach rather than relying on traditional taxonomy. In this case, our species delimitation efforts have provided new insights into patterns of fine-scale endemism in Madagascar. On a final note, there are many lineages of birds on Madagascar that are largely unstudied from an evolutionary point of view. Given several recent findings of cryptic taxa, there may be yet more species awaiting discovery in this imperiled biodiversity hotspot.

## Funding

This work was supported by the National Science Foundation (grant number DEB-1457624) awarded to SR. Funding was also provided by the Pritzker Laboratory for Molecular Systematics and Evolution at the Field Museum of Natural History, operated with support from the Pritzker Foundation.

## Acknowledgements

We are grateful to Kayleigh Kueffner for beautifully illustrating the Bernieridae. We also thank Sarah Kurtis for her work in the molecular laboratory. We gratefully acknowledge the Field Museum of Natural History, American Museum of Natural History, Mention Zoology and Animal Biodiversity at the University of Antananarivo, Burke Museum of Natural History and Culture, Louisiana State University Museum of Natural Science, and University of Kansas Biodiversity Research Institute for tissue loans.

## Data Accessibility

The Illumina short reads are available from the NCBI sequence read archive, link_TBA and Sanger sequences are available from GenBank link_TBA.

## Author Contributions

JY collected, analyzed, and interpreted the data, wrote the manuscript, and participated in conceiving and designing the study. NB collected the phenotypic data and participated in conceiving the study and interpreting the findings. MJR and SMG collected genetic samples and participated in interpreting the data and writing the manuscript. DM, CCK, and KW carried out laboratory work. SR conceived and designed the study, analyzed the phenotypic data, and participated in interpreting the data and writing the manuscript.

