## Supplemental Data for "Diversification of a cryptic radiation, a closer look at Madagascar’s recently recognized bird family"

**Table S1. Sampling of non-Bernieridae taxa including in this study**. Species names are consistent with the Clements 2018 checklist. FMNH = Field Museum of Natural History; LSUMZB = Louisiana State University Museum of Natural Science; AMNHDOT = American Museum of Natural History; BMNHC = Burke Museum of Natural History and Culture; KUBI = University of Kansas Biodiversity Research Institute.

| **Family** | **Genus** | **Species** | **Subspecies** | **Accession number** |
| --- | --- | --- | --- | --- |
| Donacobiidae | *Donacobius* | *atricapilla* | *albovittatus* | FMNH 334543 |
| Donacobiidae | *Donacobius* | *atricapilla* | *atricapilla* | FMNH 323472 |
| Locustellidae | *Bradypterus* | *alfredi* | *alfredi* | FMNH 385084 |
| Locustellidae | *Bradypterus* | *baboecala* | *centralis* | FMNH 434544 |
| Locustellidae | *Bradypterus* | *baboecala* | *tongensis* | FMNH 468159 |
| Locustellidae | *Bradypterus* | *barratti* | *cathkinensis* | FMNH 390137 |
| Locustellidae | *Bradypterus* | *brunneus* |  | FMNH 381475 |
| Locustellidae | *Bradypterus* | *brunneus* |  | FMNH 384804 |
| Locustellidae | *Bradypterus* | *carpalis* |  | FMNH 385082 |
| Locustellidae | *Bradypterus* | *cinnamomeus* | *cinnamomeus* | FMNH 385087 |
| Locustellidae | *Bradypterus* | *cinnamomeus* | *mildbreadi* | FMNH 481164 |
| Locustellidae | *Bradypterus* | *cinnamomeus* | *nyassae* | FMNH 440660 |
| Locustellidae | *Bradypterus* | *graueri* |  | FMNH 438843 |
| Locustellidae | *Bradypterus* | *lopezi* | *granti* | FMNH 447572 |
| Locustellidae | *Locustella* | *accentor* |  | LSUMZB 52679 |
| Locustellidae | *Locustella* | *amnicola* |  | BMNHC 82988 |
| Locustellidae | *Locustella* | *castanea* | *castanea* | AMNHDOT 12551 |
| Locustellidae | *Locustella* | *caudata* | *caudata* | FMNH 478821 |
| Locustellidae | *Locustella* | *caudata* | *malindangensis* | FMNH 357480 |
| Locustellidae | *Locustella* | *caudata* | *unicolor* | FMNH 472691 |
| Locustellidae | *Locustella* | *certhiola* |  | AMNHDOT 17245 |
| Locustellidae | *Locustella* | *fasciolata* |  | AMNHDOT 21951 |
| Locustellidae | *Locustella* | *fluviatilis* |  | BMNHC 82215 |
| Locustellidae | *Locustella* | *luscinioides* |  | BMNHC 72056 |
| Locustellidae | *Locustella* | *mandelli* |  | KUBI 122801 |
| Locustellidae | *Locustella* | *naevia* |  | BMNHC 64744 |
| Locustellidae | *Locustella* | *ochotensis* |  | FMNH 358393 |
| Locustellidae | *Locustella* | *tacsanowskia* |  | BMNHC 59992 |
| Locustellidae | *Megalurus* | *cruralis* |  | AMNHDOT 19773 |
| Locustellidae | *Megalurus* | *mathewsi* |  | AMNHDOT 19619 |
| Locustellidae | *Megalurus* | *palustris* | *forbesi* | FMNH 358387 |
| Locustellidae | *Megalurus* | *timoriensis* | *tweeddalei* | FMNH 472784 |
| Locustellidae | *Robsonius* | *sorsogonensis* |  | FMNH 472603 |
| Locustellidae | *Robsonius* | *thompsoni* |  | FMNH 472602 |
| Locustellidae | *Trichocichla* | *rufa* |  | KUBI 126596 |
| Acrocephalidae | *Acrocephalus* | *gracilirostris* | *jacksoni* | FMNH 434566 |
| Acrocephalidae | *Acrocephalus* | *newtoni* |  | FMNH 380001 |
| Acrocephalidae | *Acrocephalus* | *orientalis* |  | FMNH 345000 |
| Vangidae | *Vanga* | *curvirostris* |  | FMNH 384777 |

**Table S2. Bernieridae taxon sampling for this study,** with clade (*Bernieria* only), specimen identifier/s, simplified collection locality, latitude (Lat), longitude (Long), elevation, and whether the specimen was used in the family-level phylogeny. FMNH = Field Museum of Natural History; UADBA = Mention Zoologie Biologie Animale, Université d’Antananarivo; MJR = field collections of Marie-Jeanne Raherilalao held in the UADBA. Protected area locality abbreviations: MN = Monument Naturel, PHP = Paysages Harmonieux Protégé, PN= Parc National, RRN = Réserve de Ressources Naturelles, RS = Réserve Spéciale

| **Genus** | **Species** | **Clade (*Bernieria* only)** | **Specimen identifier/s** | **Simplified locality** | **Phylogeny** | **Lat** | **Long** | **Elevation** |
| --- | --- | --- | --- | --- | --- | --- | --- | --- |
| *Bernieria* | *madagascariensis* | NE (*Bernieria* sp. nov.) | MJR 1161 | PN de Masoala |  | -15.453 | 50.311 | 116 |
| *Bernieria* | *madagascariensis* | NE (*Bernieria* sp. nov.) | MJR 1181 | PN de Masoala |  | -15.453 | 50.311 | 116 |
| *Bernieria* | *madagascariensis* | NE (*Bernieria* sp. nov.) | MJR 1195 | PN de Masoala |  | -15.560 | 50.282 | 30 |
| *Bernieria* | *madagascariensis* | NE (*Bernieria* sp. nov.) | FMNH 393255 | PHP du COMATSA Sud |  | -14.543 | 49.417 | 1200 |
| *Bernieria* | *madagascariensis* | NE (*Bernieria* sp. nov.) | FMNH 356644 | Hiaraka | X | -15.483 | 49.933 | 600 |
| *Bernieria* | *madagascariensis* | SE (*Bernieria madagascariensis*) | MJR 0628; UADBA 31217 | PN de Befotaka-Midongy |  | -23.778 | 47.023 | 815 |
| *Bernieria* | *madagascariensis* | SE (*Bernieria madagascariensis*) | MJR 0670; UADBA 31255 | PN de Befotaka-Midongy |  | -23.888 | 46.897 | 1055 |
| *Bernieria* | *madagascariensis* | SE (*Bernieria madagascariensis*) | FMNH 352844 | Manantantely |  | -24.983 | 46.933 | 100 |
| *Bernieria* | *madagascariensis* | SE (*Bernieria madagascariensis*) | FMNH 345734 | PN d'Andohahela | X | -24.567 | 46.800 | 725 |
| *Bernieria* | *madagascariensis* | WS (*Bernieria inceleber*) | MJR 0541; UADBA 31029 | PN de Kirindy Mité |  | -20.788 | 44.102 | 40 |
| *Bernieria* | *madagascariensis* | WS (*Bernieria inceleber*) | MJR 0698; UADBA 31353 | RRN de Maromizaha |  | -18.976 | 48.458 | 980 |
| *Bernieria* | *madagascariensis* | WS (*Bernieria inceleber*) | MJR 0701; UADBA 31356 | RRN de Maromizaha |  | -18.976 | 48.458 | 980 |
| *Bernieria* | *madagascariensis* | WS (*Bernieria inceleber*) | MJR 0739; UADBA 31095 | RS d'Ambohitantely |  | -18.229 | 47.285 | 1425 |
| *Bernieria* | *madagascariensis* | WS (*Bernieria inceleber*) | MJR 0862; UADBA 31443 | PHP de Beanka | X | -18.062 | 44.525 | 320 |
| *Bernieria* | *madagascariensis* | WS (*Bernieria inceleber*) | MJR 0864; UADBA 31444 | PHP de Beanka |  | -18.062 | 44.525 | 320 |
| *Bernieria* | *madagascariensis* | WS (*Bernieria inceleber*) | MJR 0868; UADBA 31370 | RS d'Andranomena |  | -20.175 | 44.475 | 18 |
| *Bernieria* | *madagascariensis* | WS (*Bernieria inceleber*) | MJR 0881a; UADBA 31381 | RS d'Andranomena |  | -20.148 | 44.515 | 20 |
| *Bernieria* | *madagascariensis* | WS (*Bernieria inceleber*) | MJR 1074 | PHP de Bemanevika |  | -14.383 | 48.587 | 1517 |
| *Bernieria* | *madagascariensis* | WS (*Bernieria inceleber*) | FMNH 427373 | PHP du Corridor Forestier Ambositra-Vondrozo |  | -21.512 | 47.400 | 1075 |
| *Bernieria* | *madagascariensis* | WS (*Bernieria inceleber*) | FMNH 393260 | PHP du Corridor Forestier Ambositra-Vondrozo |  | -22.163 | 47.033 | 1600 |
| *Bernieria* | *madagascariensis* | WS (*Bernieria inceleber*) | FMNH 396156 | RS de Manongarivo |  | -13.977 | 48.417 | 785 |
| *Bernieria* | *madagascariensis* | WS (*Bernieria inceleber*) | FMNH 434671 | PN d'Ankarafantsika |  | -16.583 | 47.367 |  |
| *Bernieria* | *madagascariensis* | WS (*Bernieria inceleber*) | FMNH 479592 | PN de Ranomafana |  | -21.236 | 47.397 | 1150 |
| *Bernieria* | *madagascariensis* | SE (*Bernieria madagascariensis*) | FMNH 479595 | Forêt d'Analalava |  | -23.475 | 47.125 | 630 |
| *Bernieria* | *madagascariensis* | SE (*Bernieria madagascariensis*) | FMNH 479603 | Forêt d'Analalava |  | -22.795 | 47.188 | 590 |
| *Bernieria* | *madagascariensis* | SE (*Bernieria madagascariensis*) | FMNH 479613 | RS de Manombo |  | -23.030 | 47.710 | 60 |
| *Bernieria* | *madagascariensis* | WS (*Bernieria inceleber*) | FMNH 479620 | Forêt d'Ankazomivady |  | -20.773 | 47.183 | 1710 |
| **Genus** | **Species** | **Clade (*Bernieria* only)** | **Specimen identifier/s** | **Simplified locality** | **Phylogeny** | **Lat** | **Long** | **Elevation** |
| *Bernieria* | *madagascariensis* | WS (*Bernieria inceleber*) | FMNH 479624 | Forêt de Mahatsinjo (Tsinjoarivo) |  | -19.684 | 47.755 | 1600 |
| *Bernieria* | *madagascariensis* | WS (*Bernieria inceleber*) | FMNH 427371 | MN Forèt Sacrée d'Alandraza Analavelo |  | -22.642 | 44.167 | 1250 |
| *Bernieria* | *madagascariensis* | WS (*Bernieria inceleber*) | FMNH 431205 | PN de Bemaraha |  | -19.132 | 44.800 | 100 |
| *Bernieria* | *madagascariensis* | WS (*Bernieria inceleber*) | FMNH 436428 | PN de Mikea |  | -22.767 | 43.517 | 80 |
| *Bernieria* | *madagascariensis* | WS (*Bernieria inceleber*) | FMNH 436505 | PN de Namoroka |  | -16.433 | 45.400 | 120 |
| *Bernieria* | *madagascariensis* | SE (*Bernieria madagascariensis*) | FMNH 352861 | Forêt de Marovony |  | -24.100 | 47.367 | 50 |
| *Bernieria* | *madagascariensis* | SE (*Bernieria madagascariensis*) | FMNH 363819 | PN d'Andringitra |  | -22.222 | 47.017 | 720 |
| *Bernieria* | *madagascariensis* | WS (*Bernieria inceleber*) | FMNH 393258 | PHP du Corridor Forestier Ambositra-Vondrozo |  | -22.149 | 47.017 | 1300 |
| *Bernieria* | *madagascariensis* | WS (*Bernieria inceleber*) | FMNH 393259 | PHP du Corridor Forestier Ambositra-Vondrozo |  | -22.149 | 47.017 | 1300 |
| *Bernieria* | *madagascariensis* | WS (*Bernieria inceleber*) | FMNH 393263 | RS d'Ambohijanahary |  | -18.262 | 45.417 | 1150 |
| *Bernieria* | *madagascariensis* | WS (*Bernieria inceleber*) | FMNH 396157 | RS de Manongarivo |  | -13.999 | 48.417 | 1240 |
| *Bernieria* | *madagascariensis* | NE (*Bernieria* sp. nov.) | FMNH 431201 | PN de Marojejy |  | -14.437 | 49.617 | 1175 |
| *Crossleyia* | *xanthophrys* |  | MJR 0391; UADBA 30915 | PHP du Complexe Anjozorobe-Angavo |  | -18.425 | 47.953 | 1300 |
| *Crossleyia* | *xanthophrys* |  | MJR 0405; UADBA 31280 | PHP du Complexe Anjozorobe-Angavo | X | -18.473 | 47.960 | 1325 |
| *Cryptosylvicola* | *randrianasoloi* |  | FMNH 360058 | RRN de Maromizaha |  | -18.983 | 48.450 | 1100 |
| *Cryptosylvicola* | *randrianasoloi* |  | FMNH 360059 | RRN de Maromizaha |  | -18.983 | 48.450 | 1100 |
| *Cryptosylvicola* | *randrianasoloi* |  | FMNH 384748 | PN de Marojejy |  | -14.447 | 49.733 | 1875 |
| *Cryptosylvicola* | *randrianasoloi* |  | FMNH 396198 | RS de Manongarivo | X | -14.022 | 48.417 | 1600 |
| *Hartertula* | *flavoviridis* |  | MJR 0294; UADBA 47231 | RS de Marotandrano |  | -16.285 | 48.815 | 850 |
| *Hartertula* | *flavoviridis* |  | MJR 0590; UADBA 31183 | PN de Befotaka-Midongy | X | -23.737 | 47.023 | 835 |
| *Hartertula* | *flavoviridis* |  | MJR 0714; UADBA 31075 | RRN de Maromizaha |  | -18.976 | 48.458 |  |
| *Oxylabes* | *madagascariensis* | | MJR 0280; UADBA 47241 | RS de Manongarivo |  | -16.280 | 48.802 | 950 |
| *Oxylabes* | *madagascariensis* | | MJR 0365; UADBA 47982 | PHP du Complexe Anjozorobe-Angavo |  | -18.422 | 47.937 | 1250 |
| *Oxylabes* | *madagascariensis* | | MJR 0647; UADBA 31236 | PN de Befotaka-Midongy |  | -23.837 | 46.960 | 1026 |
| *Oxylabes* | *madagascariensis* | | MJR 0703; UADBA 31069 | RRN de Maromizaha |  | -18.976 | 48.458 | 980 |
| *Oxylabes* | *madagascariensis* | | MJR 0815; UADBA 31166 | Ambohimanarivo |  | -18.806 | 48.361 | 1006 |
| *Oxylabes* | *madagascariensis* | | MJR 0892; UADBA 31400 | RRH du Corridor Ankeniheny-Zahamena |  | -19.044 | 48.349 | 997 |
| **Genus** | **Species** | **Clade (*Bernieria* only)** | **Specimen identifier/s** | **Simplified locality** | **Phylogeny** | **Lat** | **Long** | **Elevation** |
| *Oxylabes* | *madagascariensis* |  | MJR 1202 | PN de Masoala |  | -15.560 | 50.282 | 30 |
| *Oxylabes* | *madagascariensis* |  | FMNH 352915 | Forêt de Marovony | X | -24.100 | 47.367 | 50 |
| *Oxylabes* | *madagascariensis* |  | MJR 0758; UADBA 31112 | Ambatovy-Analamay |  | -18.799 | 48.323 |  |
| *Oxylabes* | *madagascariensis* |  | MJR 1235 | PN de Masoala |  | -15.569 | 50.003 | 610 |
| *Randia* | *pseudozosterops* |  | FMNH 479641 | PN de Ranomafana | X | -21.236 | 47.397 | 1150 |
| *Thamnornis* | *chloropetoides* |  | MJR 0323; UADBA 47401 | Forêt de Tongaenoro |  | -24.737 | 44.030 | 120 |
| *Thamnornis* | *chloropetoides* |  | MJR 0341; UADBA 47752 | PHP de Nord-Ifotaka |  | -24.765 | 46.153 | 110 |
| *Thamnornis* | *chloropetoides* |  | MJR 0554; UADBA 31041 | PN de Kirindy Mité |  | -20.942 | 43.872 | 10 |
| *Thamnornis* | *chloropetoides* |  | MJR 0916; UADBA 33497 | PN de Tsimanampesotse |  | -24.027 | 43.737 | 19 |
| *Thamnornis* | *chloropetoides* |  | MJR 0955; UADBA 33590 | PN de Mikea | X | -22.510 | 43.295 | 17 |
| *Xanthomixis* | *apperti* |  | FMNH 393162 | MN Forèt Sacrée d'Alandraza Analavelo |  | -22.678 | 44.183 | 1050 |
| *Xanthomixis* | *apperti* |  | FMNH 427363 | MN Forèt Sacrée d'Alandraza Analavelo | X | -22.642 | 44.167 | 1250 |
| *Xanthomixis* | *cinereiceps* |  | MJR 0424; UADBA 30940 | PHP du Complexe Anjozorobe-Angavo |  | -18.522 | 47.973 | 1280 |
| *Xanthomixis* | *cinereiceps* |  | MJR 0651; UADBA 31240 | PN de Befotaka-Midongy |  | -23.837 | 46.960 | 1026 |
| *Xanthomixis* | *cinereiceps* |  | MJR 0695; UADBA 31067 | RRN de Maromizaha |  | -18.976 | 48.458 | 980 |
| *Xanthomixis* | *cinereiceps* |  | MJR 1090 | PHP de Bemanevika | X | -14.383 | 48.587 | 1517 |
| *Xanthomixis* | *tenebrosa* |  | FMNH 393278 | PHP du COMATSA Sud | X | -14.538 | 49.433 | 875 |
| *Xanthomixis* | *zosterops* |  | MJR 0591 | PN de Befotaka-Midongy |  | -23.737 | 47.023 | 835 |
| *Xanthomixis* | *zosterops* |  | MJR 0630 | PN de Befotaka-Midongy |  | -23.778 | 47.023 | 815 |
| *Xanthomixis* | *zosterops* |  | MJR 0631 | PN de Befotaka-Midongy |  | -23.778 | 47.023 | 815 |
| *Xanthomixis* | *zosterops* |  | MJR 0657 | PN de Befotaka-Midongy |  | -23.888 | 46.897 | 1055 |
| *Xanthomixis* | *zosterops* |  | MJR 0776 | Ambatovy-Analamay |  | -18.793 | 48.334 | 1105 |
| *Xanthomixis* | *zosterops* |  | MJR 1159 | PN de Masoala |  | -15.453 | 50.311 | 116 |
| *Xanthomixis* | *zosterops* |  | MJR 1196 | PN de Masoala |  | -15.560 | 50.282 | 30 |
| *Xanthomixis* | *zosterops* |  | MJR 1208 | PN de Masoala |  | -15.560 | 50.282 | 30 |
| *Xanthomixis* | *zosterops* |  | MJR 1240 | PN de Masoala |  | -15.729 | 49.964 | 15 |
| *Xanthomixis* | *zosterops* |  | MJR 1241 | PN de Masoala |  | -15.729 | 49.964 | 15 |
| *Xanthomixis* | *zosterops* |  | FMNH 431197 | PN de Marojejy |  | -14.427 | 49.600 | 810 |
| *Xanthomixis* | *zosterops* |  | FMNH 427376 | MN Forèt Sacrée d'Alandraza Analavelo | X | -22.642 | 44.167 | 1250 |
| *Xanthomixis* | *zosterops* |  | FMNH 438696 | PN de Befotaka-Midongy |  | -23.835 | 46.950 | 875 |
| *Xanthomixis* | *zosterops* |  | FMNH 438698 | PN de Befotaka-Midongy |  | -23.835 | 46.950 | 875 |
| **Genus** | **Species** | **Clade (*Bernieria* only)** | **Specimen identifier/s** | **Simplified locality** | **Phylogeny** | **Lat** | **Long** | **Elevation** |
| *Xanthomixis* | *zosterops* |  | FMNH 345744 | Forêt d'Analalava |  | -24.217 | 47.317 | 40 |
| *Xanthomixis* | *zosterops* |  | FMNH 345747 | PN d'Andohahela |  | -24.567 | 46.800 | 725 |
| *Xanthomixis* | *zosterops* |  | FMNH 345749 | PN d'Andohahela |  | -24.567 | 46.800 | 725 |
| *Xanthomixis* | *zosterops* |  | FMNH 345752 | PN d'Andohahela |  | -24.567 | 46.800 | 725 |
| *Xanthomixis* | *zosterops* |  | FMNH 427374 | PN de Ranomafana |  | -21.290 | 47.433 | 1025 |
| *Xanthomixis* | *zosterops* |  | FMNH 393137 | Forêt d'Ankazomivady |  | -20.775 | 47.167 | 1675 |

**Evaluating the presence of cryptic species within Bernieridae**

We examined each of the 11 species within the Bernieridae family for evidence of cryptic species-level diversity. We did this because other avian lineages in Madagascar have recently been shown to contain cryptic species (Younger et al. In review; Younger et al. 2018), and a previous study of Bernieridae suggested it may be the case for this family also (Block 2012). In total, we sequenced 91 Bernieridae individuals for UCEs following the protocol in the main text. For each species we included representative individuals from all key biogeographic regions inhabited, as far as possible given the availability of tissues. For some species only a small number of samples are included because these species have limited geographic ranges. We also used the Block (2012) mitochondrial study as a preliminary assessment of potential divergences to lay the foundation for this more extensive study using thousands of nuclear loci. Table S1 has the full details for all these individuals, and a maximum likelihood phylogeny for all the sequenced individuals is shown in Figure S1.

We followed these steps to assess species limits:

1. Does the species have a restricted range and is therefore unlikely to support cryptic species? Do we have insufficient sampling to assess the presence of distinct lineages? If YES to either of these questions, then do not analyze species limits.
2. Does the species show geographically distinct lineages in the UCE phylogeny with high levels of bootstrap support (>90% for reciprocally monophyletic groups)?
3. Are the lineages also supported as distinct with minimal admixture in genetic clustering analyses using SNP data?

**Figure S1 (next page). UCE phylogeny for all 130 taxa included in this study**, with all bootstrap supports and accession numbers. Maximum-likelihood tree of 4317 concatenated UCE loci; 3,912,738 bp).

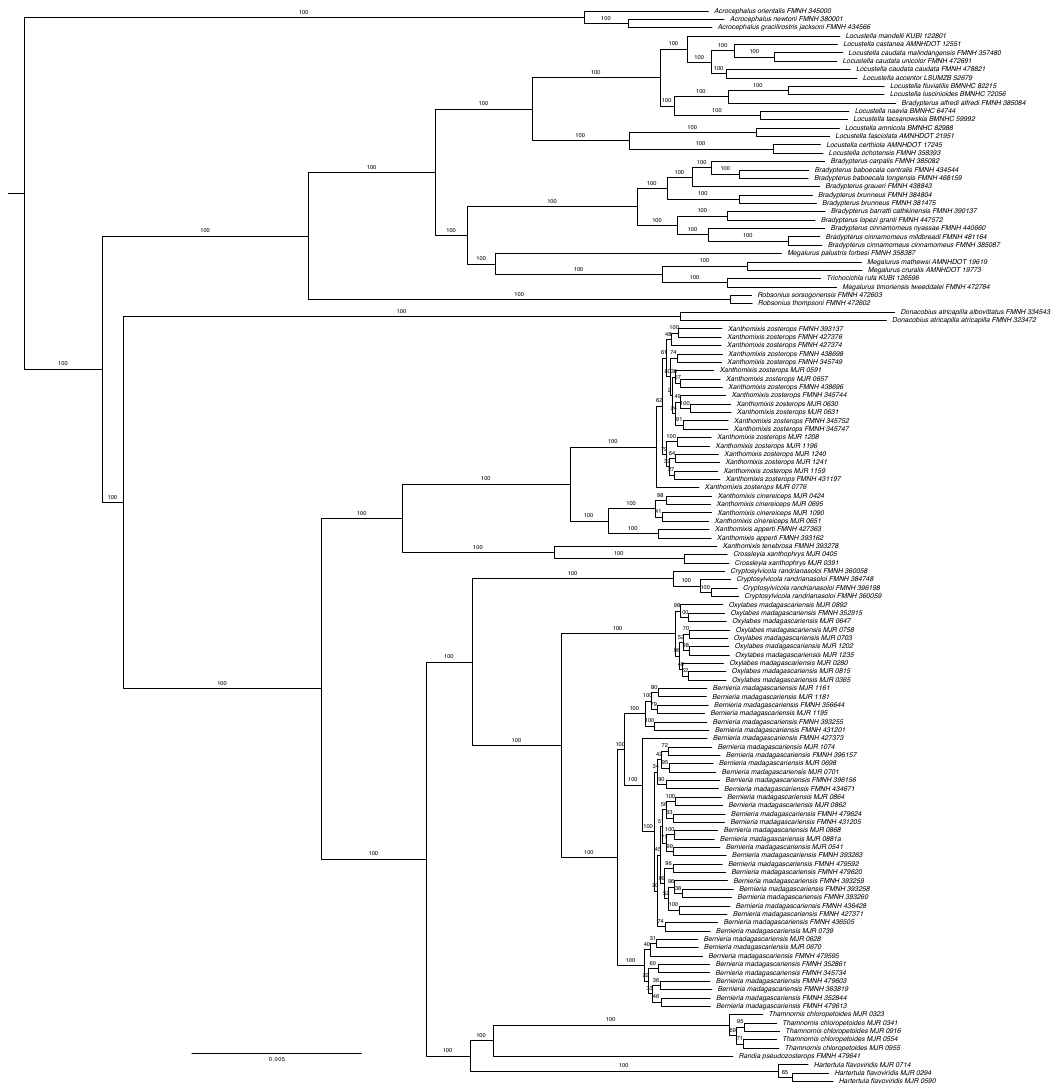

***Bernieria madagascariensis***: Twenty individuals were initially sequenced for UCEs. Within the UCE phylogeny, there were three clades with 100% bootstrap support (Figure S2). These clades were correlated with geography; one was found in the northeast, one in the southeast, and the other was widespread in western Madagascar and the central portions of the eastern forest.

We called SNPs on the 20 individuals (3576 unlinked SNPs, mean 69X coverage) and carried out a Structure (Pritchard et al. 2000) clustering analysis to determine the most likely number of genetic clusters in the dataset, and the proportion of assignment of individuals to those clusters. The optimal number of clusters was *K* = 3 based on the maximum posterior log probability. The assignment of individuals to three clusters (Figure S3) was entirely consistent with the three phylogenetic lineages, with the exception of one apparently hybrid individual (FMNH 427373). All other individuals had 100% assignment to their genetic clusters.

Overall, we recovered distinct phylogenetic lineages with 100% bootstrap support, the same groupings based on genetic clustering analyses, and no evidence of admixture except for one hybrid individual. It is possible that these three lineages represent distinct species; we therefore increased our sample size for *Bernieria* to 39 individuals to allow for better determination of geographic boundaries of these clades and conducted further analyses – please refer to the main manuscript for details.

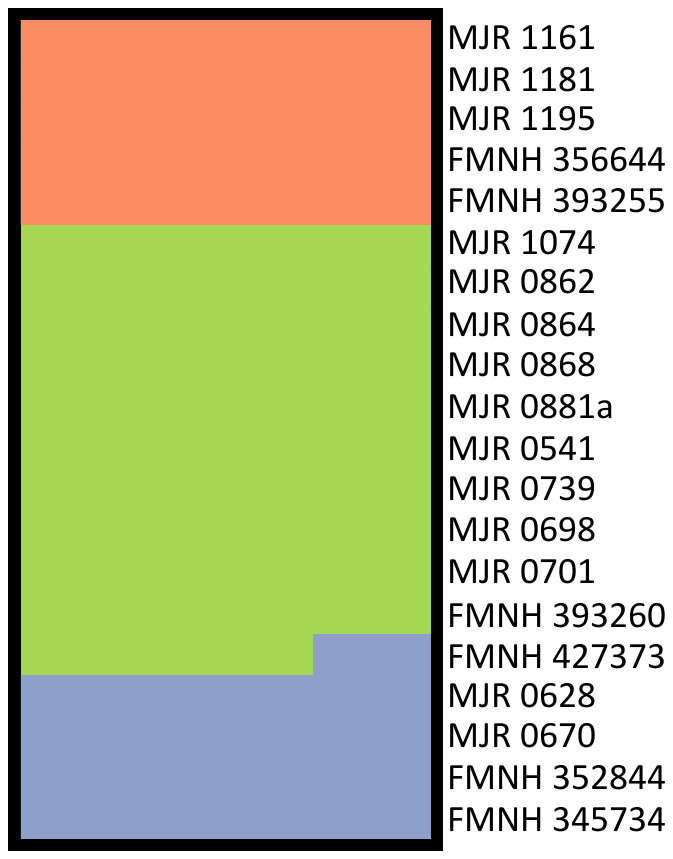

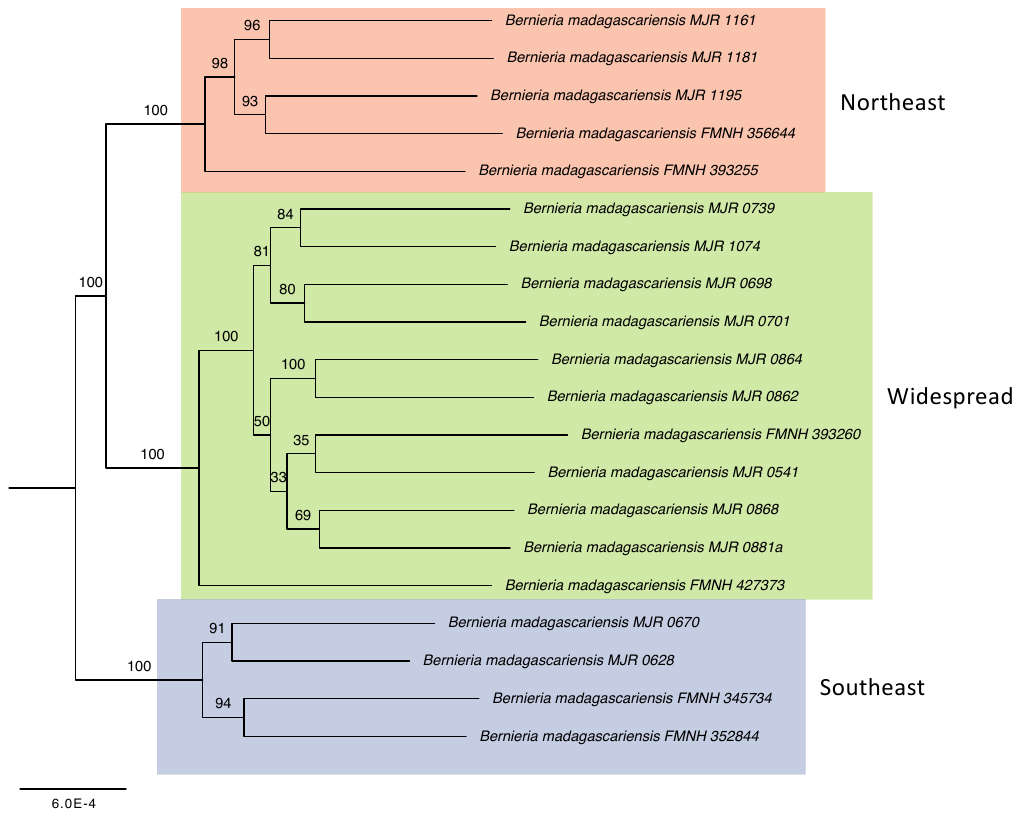

**Figure S2. *Bernieria* UCE phylogeny** (maximum-likelihood tree of 4184 concatenated UCE loci; 3,377,916 bp).

**Figure S3. Structure plot of *Bernieria* specimens, for *K* = 3.**

likelihood tree of 4184 concatenated UCE loci; 3,377,916 bp).

***Crossleyia xanthophrys***: Only two individuals were available for UCE sequencing. A preliminary study of mitochondrial DNA of eight individuals found no evidence for distinct lineages at the 2% divergence level (Block 2012).

***Cryptosylvicola randriansoloi***: Four individuals were sequenced for UCEs (Figure S5). The UCE phylogeny does not show geographically distinct, well-supported lineages (Figure S4).

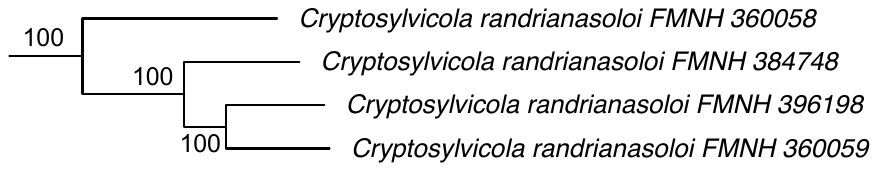

**Figure S4. *Cryptosylvicola randriansoloi* clade from UCE phylogeny** (full phylogeny shown in Figure S1).

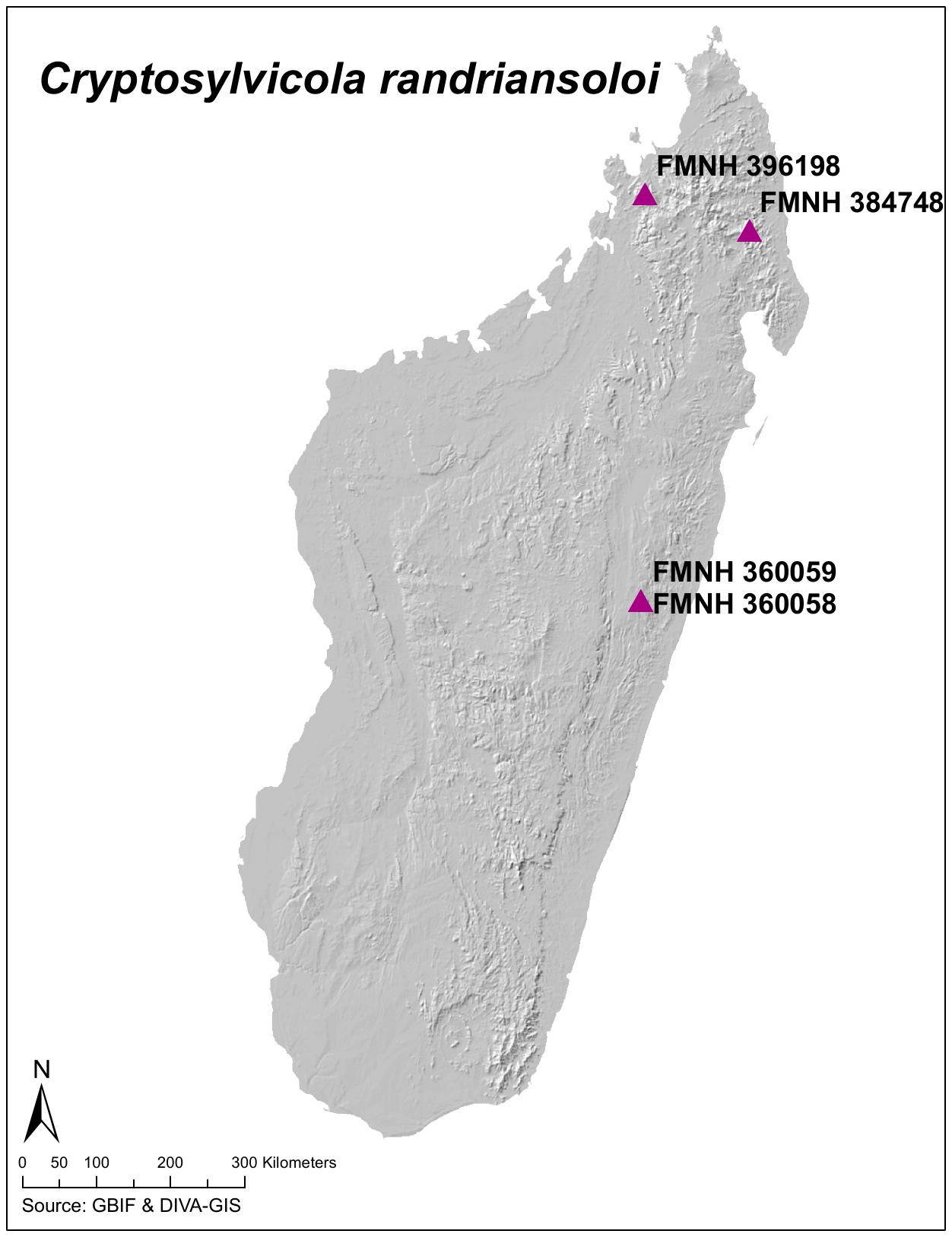

**Figure S5. Map of *Cryptosylvicola randriansoloi* specimens used in this study.**

***Hartertula flavoviridis***: Three individuals were sequenced for UCEs, from throughout the eastern humid forest (Figure S7). The UCE phylogeny does not show geographically distinct, well-supported lineages (Figure S6).

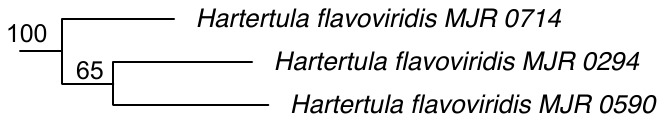

**Figure S6. *Hartertula flavoviridis* clade from UCE phylogeny** (full phylogeny shown in Figure S1).

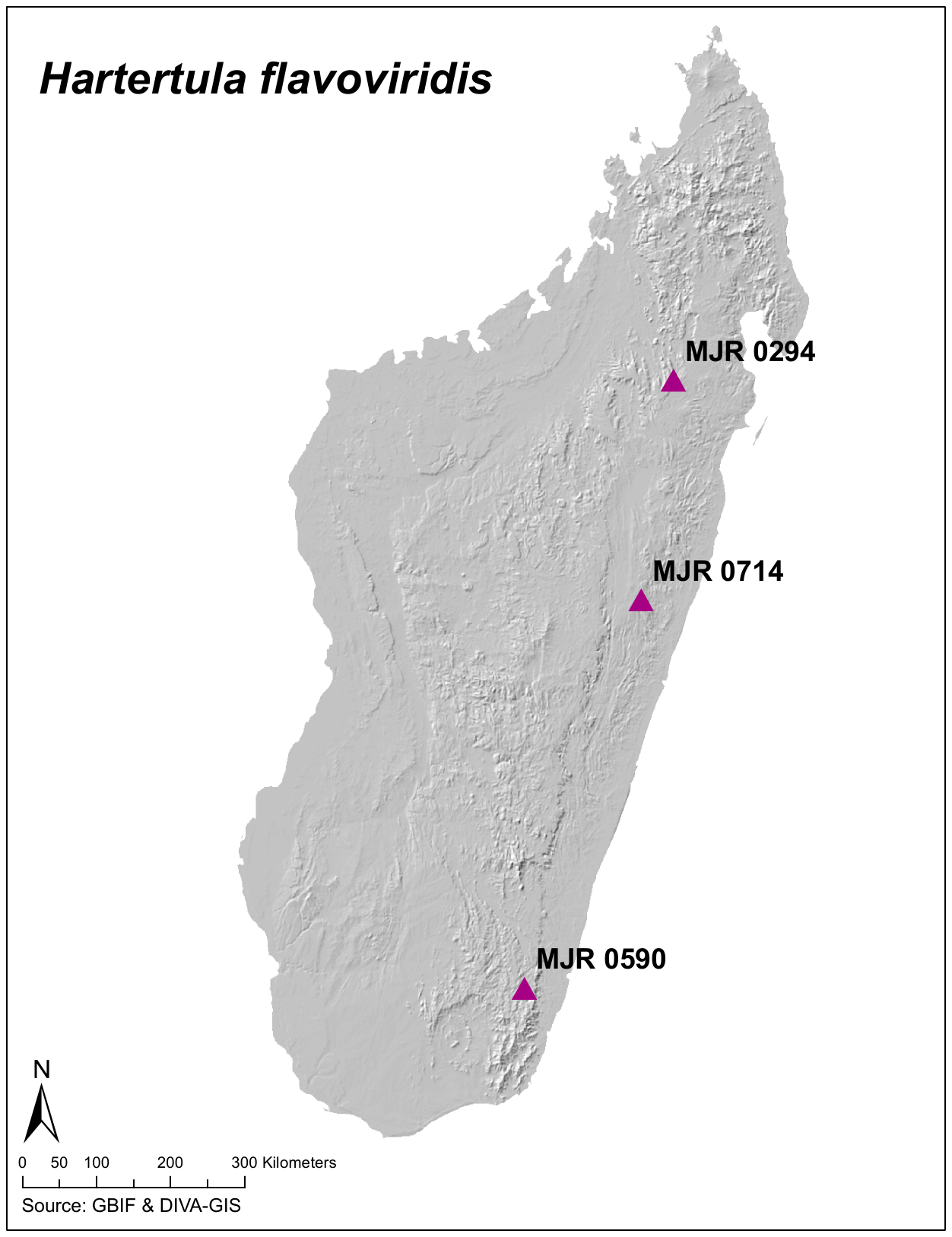

**Figure S7. Map of *Hartertula flavoviridis* specimens used in this study.**

***Oxylabes madagascariensis***: Ten individuals were sequenced for UCEs (Figure S10). Within the UCE phylogeny, there were two potentially distinct lineages, each with 98% bootstrap support (Figure S8). These clades are geographically separated by only 14 km in the central eastern forest (Figure S10); one clade is composed of individuals from the central and northern zones of the eastern humid forest, the other consists of individuals from the central and southern areas of the eastern forests. We called SNPs on the 10 individuals (3239 unlinked SNPs, mean coverage 62X) and carried out a Structure (Pritchard et al. 2000) clustering analysis to determine the most likely number of genetic clusters in the dataset, and the proportion of assignment of individuals to those clusters. The maximum posterior log probability was achieved at *K* = 2 (-31534), however this was only a minor improvement over *K =* 1 (-31275), therefore either one or two genetic clusters may be biologically plausible (Figure S11). For *K* = 2, the two southernmost individuals were assigned to one cluster, and the remainder (central and northern) to the other (Figure S9). An individual from the Central Highlands (MJR 0892) that fell within the southern phylogenetic lineage was assigned predominantly to the northern cluster in the Structure analysis. Overall, these findings do not provide convincing evidence of species-level divergences within *O. madagascariensis*, but are suggestive of some genetic divergence over large distances, as expected for non-dispersive forest taxa.

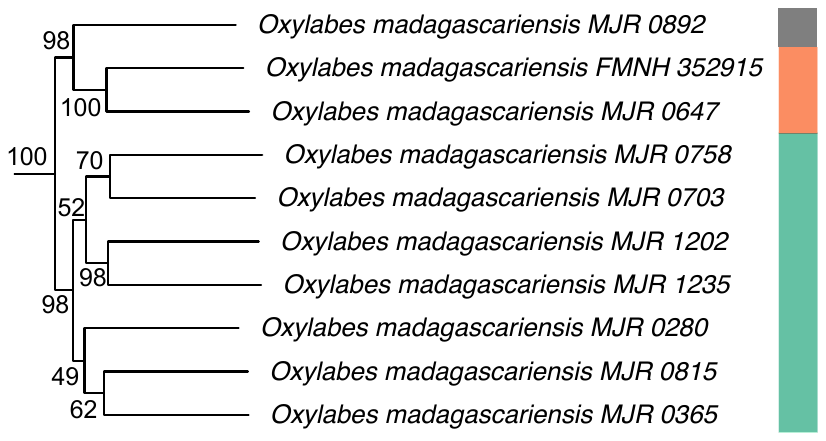

**Figure S8. *Oxylabes madagascariensis* clade from UCE phylogeny** (full phylogeny shown in Figure S1). Colors match the northern (green) and southern (orange) clades/genetic clusters, with the undefinitively assigned individual (MJR0982) shown in black.

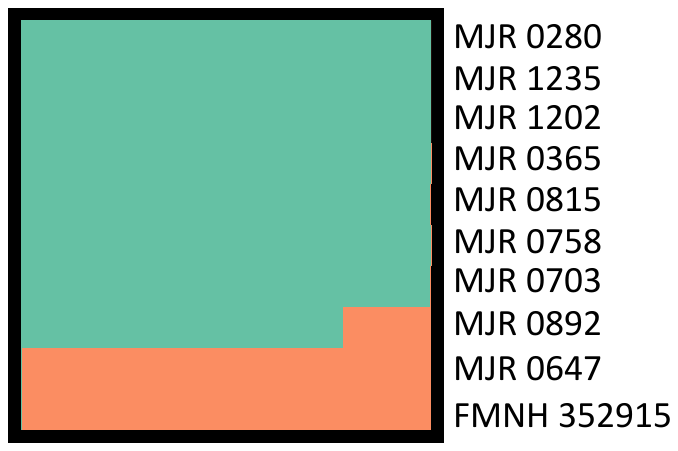

**Figure S9. Structure plot of *Oxylabes madagascariensis* specimens, for *K = 2*.**

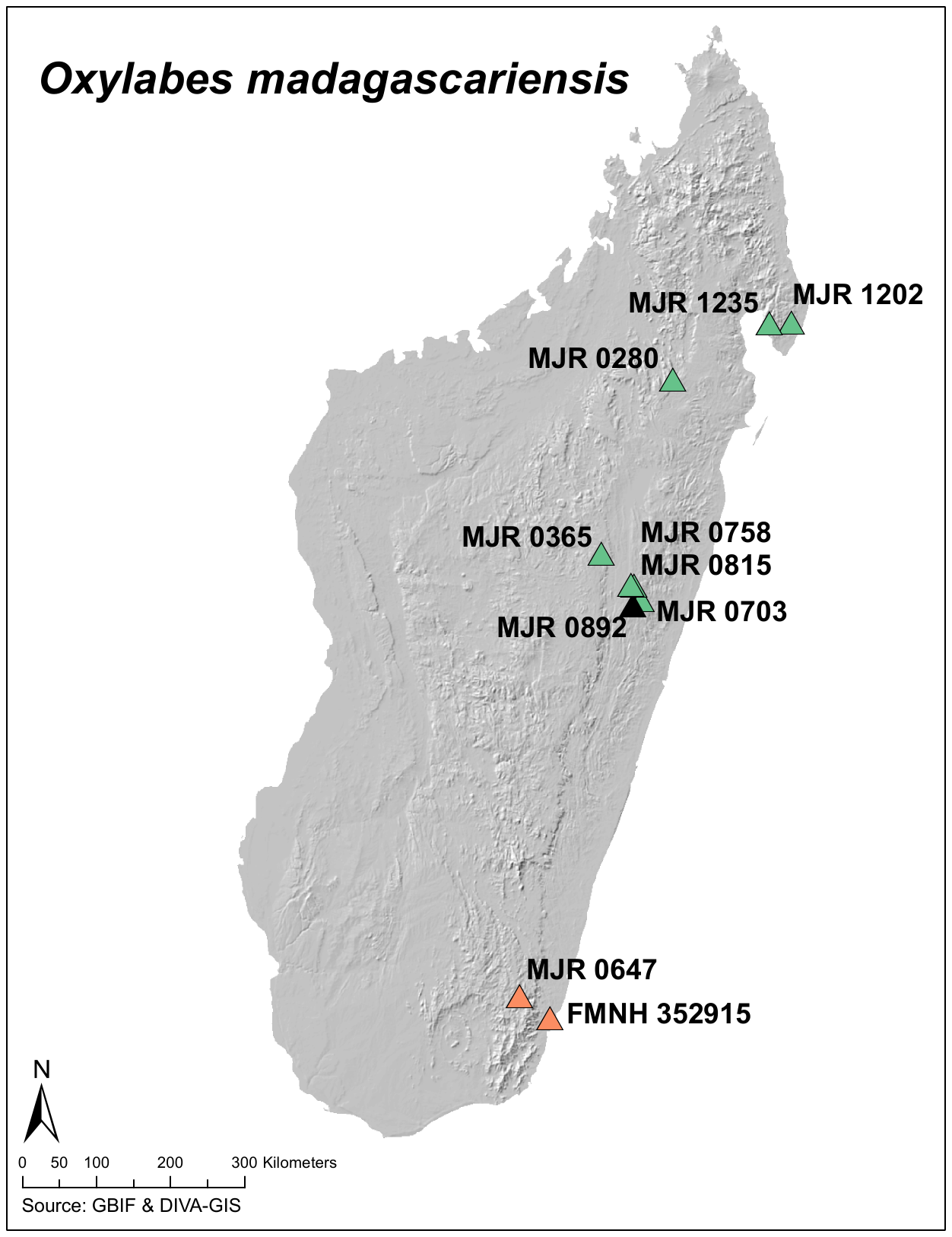

**Figure S10. Map of *Oxylabes madagascariensis* specimens used in this study**. Colors match the northern and southern clades/genetic clusters, with the non-definitively assigned individual (MJR0982) shown in black.

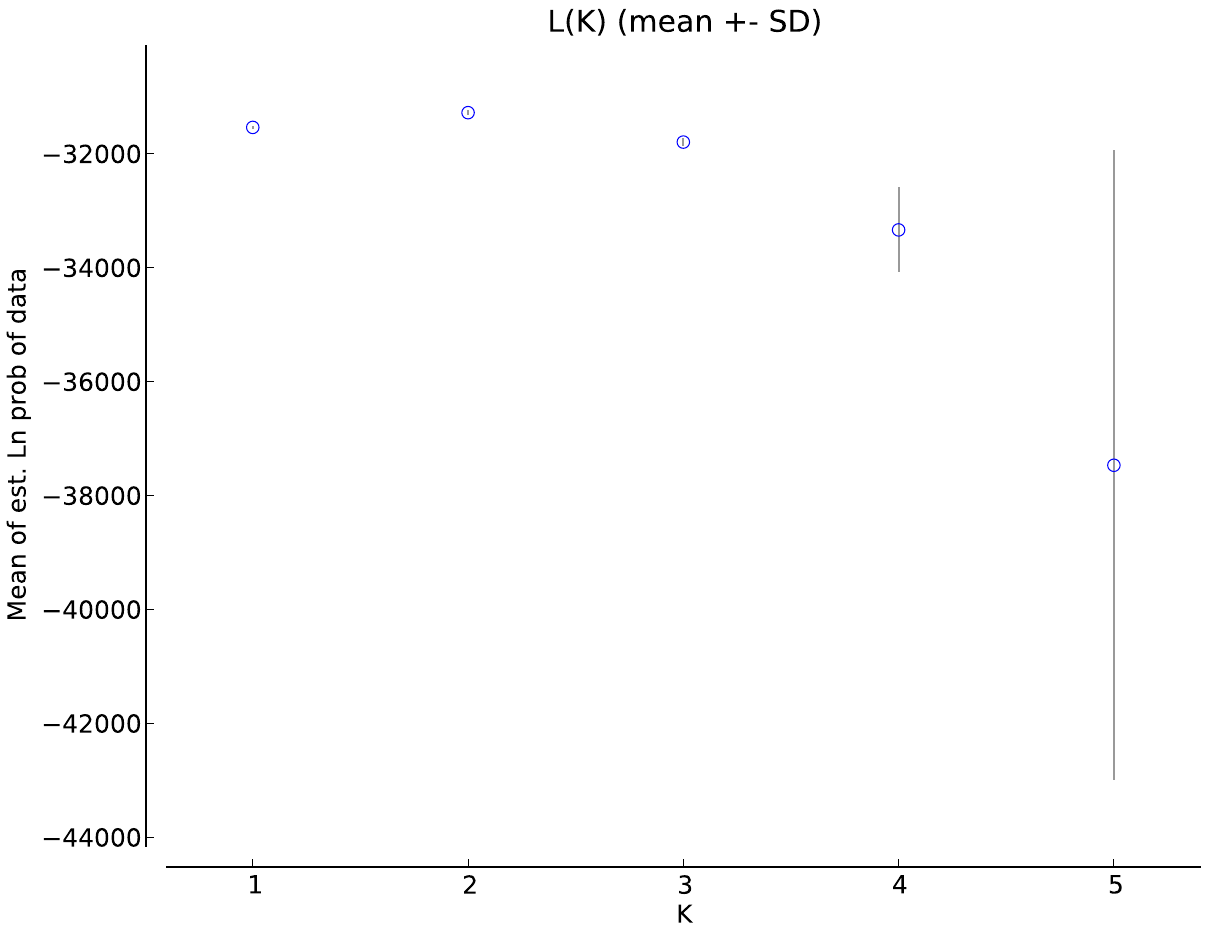

**Figure S11. Estimated log probability of the data Pr(*X*|*K*) for each value of *K* for *Oxylabes madagascariensis***, averaged over ten independent Structure runs.

***Randia pseudozosterops***: This is a small, canopy-living species rarely caught in field surveys, and only one specimen was available for sequencing. However, *R. pseudozosterops* is distributed throughout the eastern humid forest (Goodman and Raherilalao 2013) and further phylogeographic study is warranted if additional genetic samples can be obtained in the future.

***Thamnornis chloropetoides***: Five individuals were sequenced for UCEs. The species has a relatively broad zone of occurrence from the southeast across the south to the southwest. Specimens from across this area were included in our analysis, and the UCE phylogeny does not show any of these populations to be phylogenetically divergent.

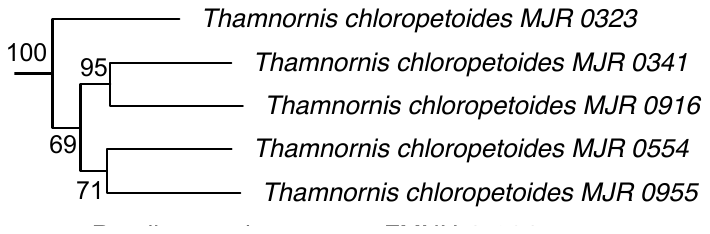

**Figure S12. *Thamnornis chloropetoides* clade from UCE phylogeny** (full phylogeny shown in Figure S1).

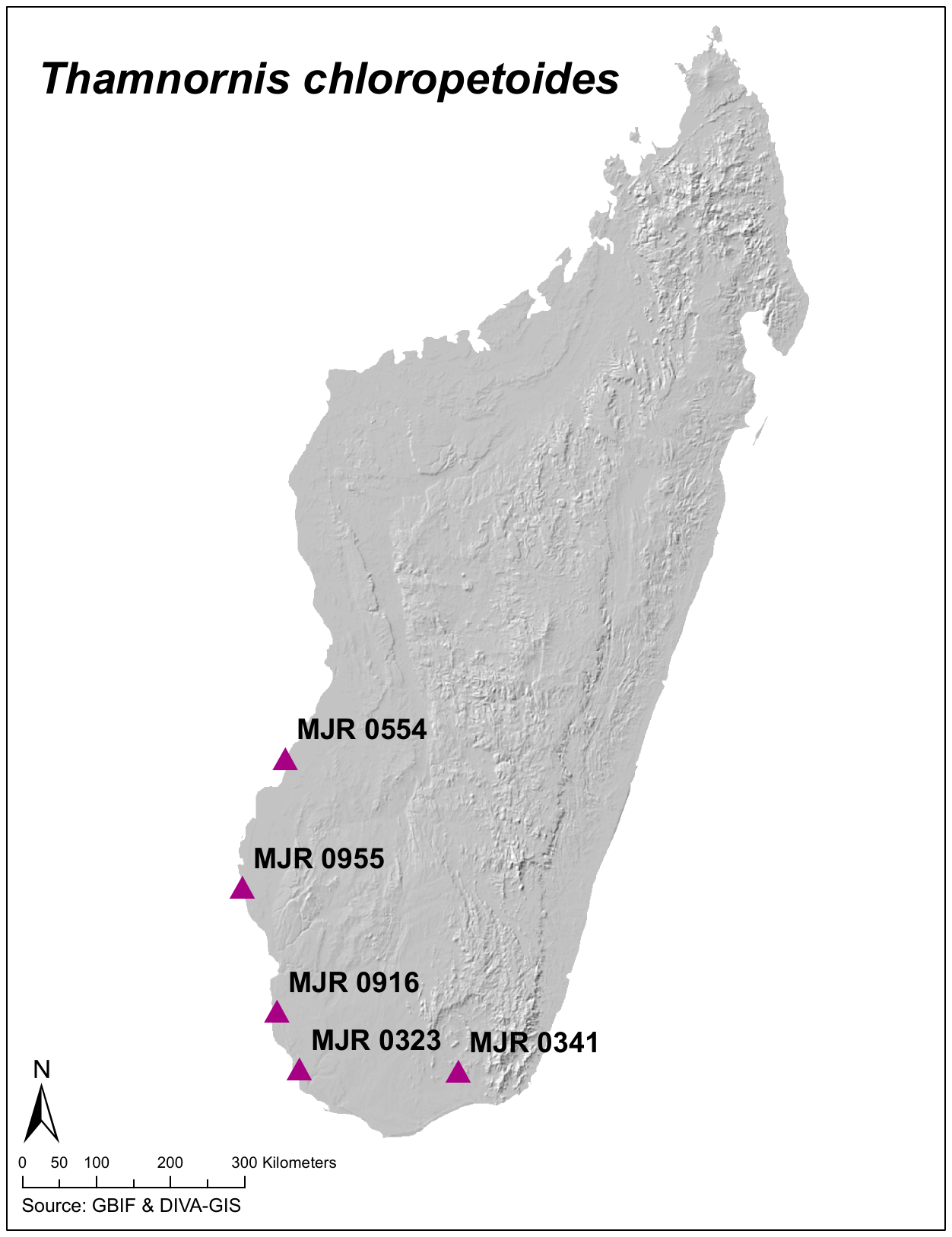

**Figure S13. Map of *Thamnornis chloropetoides* specimens used in this study.**

***Xanthomixis apperti***: This species has a relatively small distribution in south central and southwestern Madagascar (Goodman and Raherilalao 2013) that is unlikely to support distinct genetic lineages. The species is also considered reclusive with very few records (Goodman and Raherilalao 2013), and only two individuals were available for UCE sequencing.

***Xanthomixis cinereiceps***: Four individuals were sequenced for UCEs, from throughout the eastern humid forest (Figure S15). The UCE phylogeny does not show geographically distinct, well-supported lineages (Figure S14).

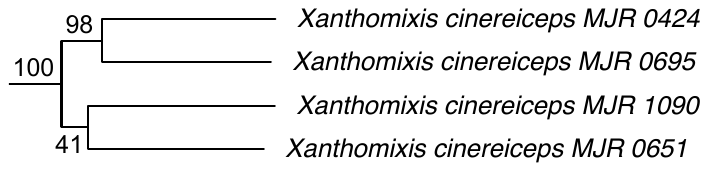

**Figure S14. *Xanthomixis cinereiceps* clade from UCE phylogeny** (full phylogeny shown in Figure S1).

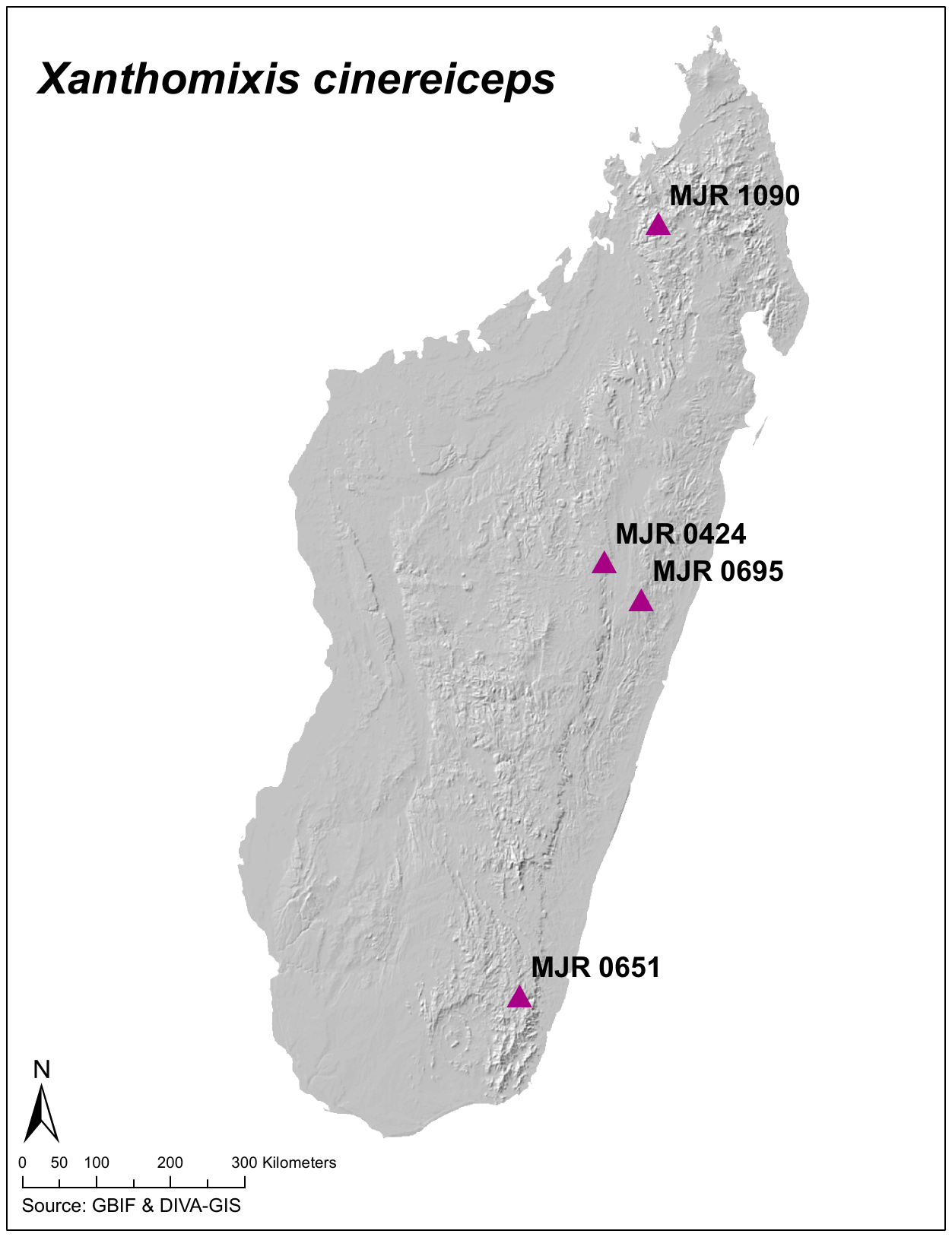

**Figure S15. Map of *Xanthomixis cinereiceps* specimens used in this study.**

***Xanthomixis tenebrosa***: Only one tissue specimen available; this species is known from only a small number of records (Goodman and Raherilalao 2013). *Xanthomixis tenebrosa* has two distinct areas of occurrence in the eastern humid forests (Goodman and Raherilalao 2013), therefore future investigation of genetic populations is warranted if genetic samples can be obtained.

***Xanthomixis zosterops***: Twenty individuals were sequenced for UCEs (Figure S16). Within the UCE phylogeny, there were three potentially distinct lineages; two of these had bootstrap supports of 90% and 84% respectively, and the third contained only a single taxon (Figure S17). These clades were loosely correlated with geography; one (90% supported) is composed of individuals from northeast Madagascar, the other (84% supported) consists of individuals from southeast and southwest Madagascar, and the most divergent individual was from the central eastern humid forest. We called SNPs on the 20 individuals (2668 unlinked SNPs, mean coverage 119X) and carried out a Structure (Pritchard et al. 2000) clustering analysis to determine the most likely number of genetic clusters in the dataset, and the proportion of assignment of *X. zosterops* individuals to those clusters. The optimal number of clusters was *K* = 3, based on both the maximum posterior log probability and the rate of change in log probability (deltaK, Evanno method (Evanno et al. 2005)). The assignment of individuals to these three clusters was inconsistent with the three phylogenetic lineages (Figure S18). The northeastern phylogenetic lineage appears as a distinct genetic cluster (green in Figure S18), and a second southeastern cluster is also apparent (orange in Figure S18). However, an individual from the southwest (FMNH 427376) that fell within the southern phylogenetic lineage appears genetically divergent in the Structure analysis, and the most divergent individual in the phylogenetic analysis (MJR 0776) appears to be an admixed individual in the clustering analysis. Several individuals appear to be of admixed ancestry, suggesting interbreeding among genetic populations. Overall, our findings suggest population genetic structure rather than species-level divergences. However, a previous phylogeographic study of *X. zosterops* found an intriguing pattern; deep mitochondrial divergences, yet near panmixia based on microsatellites (Block et al. 2015). These interesting phylogeographic patterns warrant further investigation with more extensive sampling.

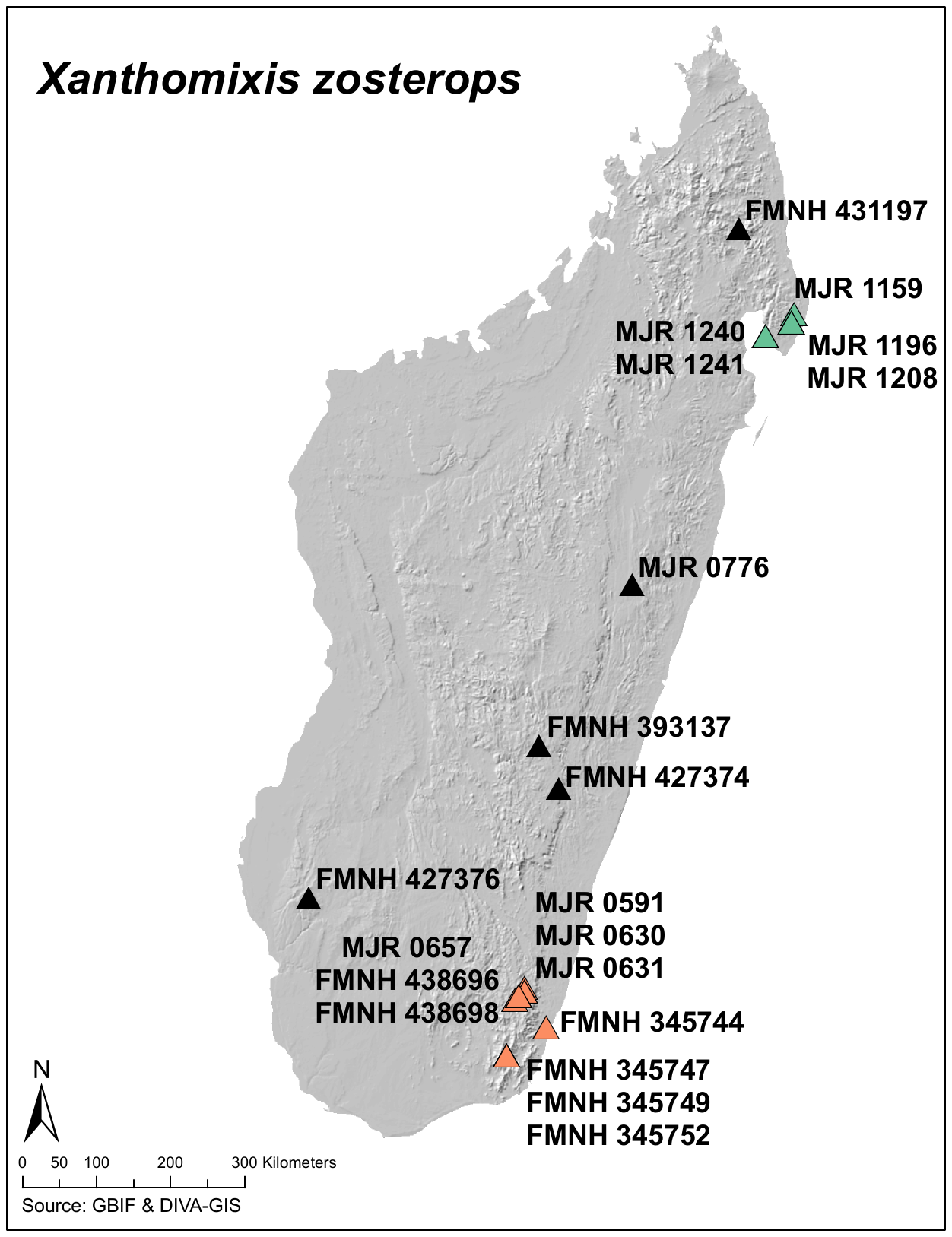

**Figure S16. Map of *Xanthomixis zosterops* specimens used in this study.** Orange individuals belong to the southeastern clade and genetic cluster, green indicates the northeastern clade and genetic cluster, black indicates admixed individuals, or different phylogenetic clades compared to Structure clusters.

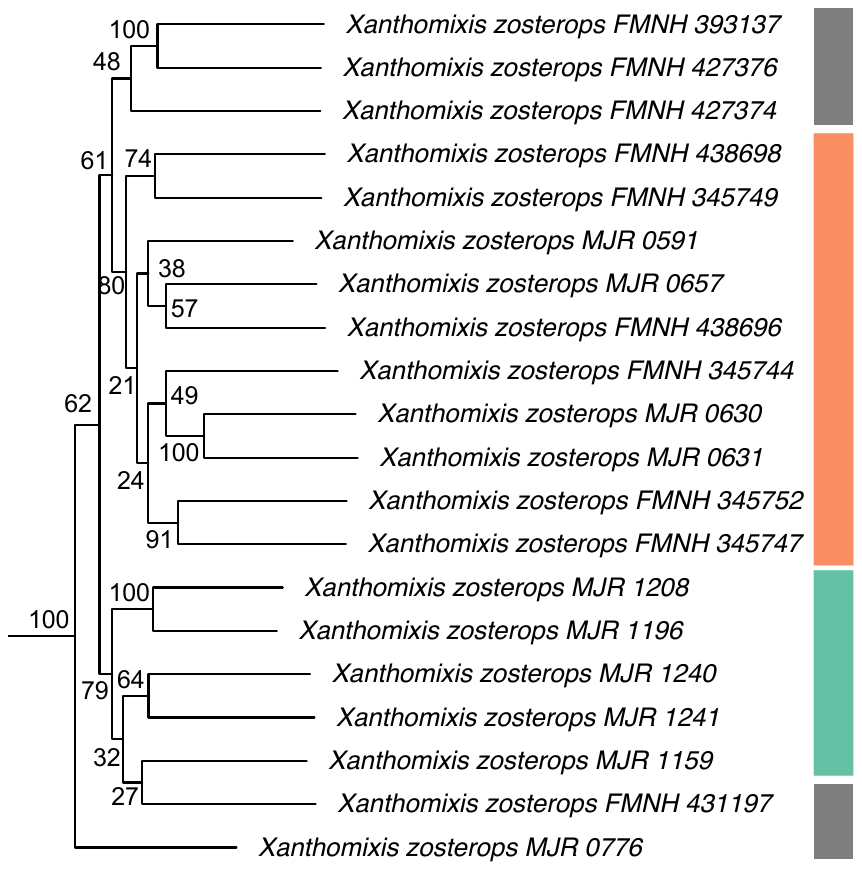

**Figure S17. *Xanthomixis zosterops* clade from UCE phylogeny** (full phylogeny shown in Figure S1). Colors correspond to those in the sampling map – orange individuals are members of the southern clade and Structure cluster, green individuals are members of the northern clade and Structure cluster, black indicates admixed individuals, or different membership in the phylogenetic compared to clustering analyses.

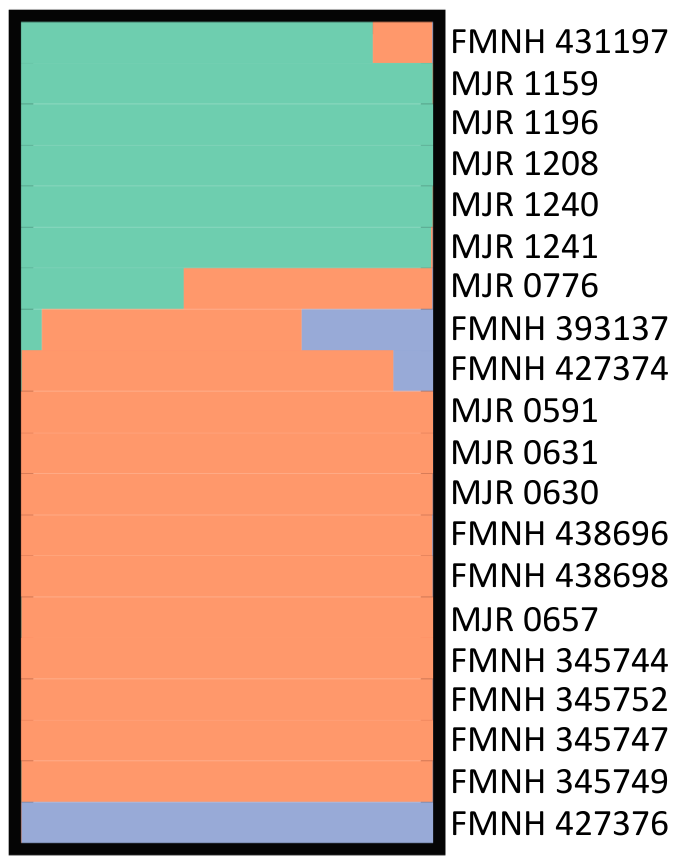

**Figure S18. *Xanthomixis zosterops* Structure plot for *K* = 3.**

**Additional supplementary figures & tables**

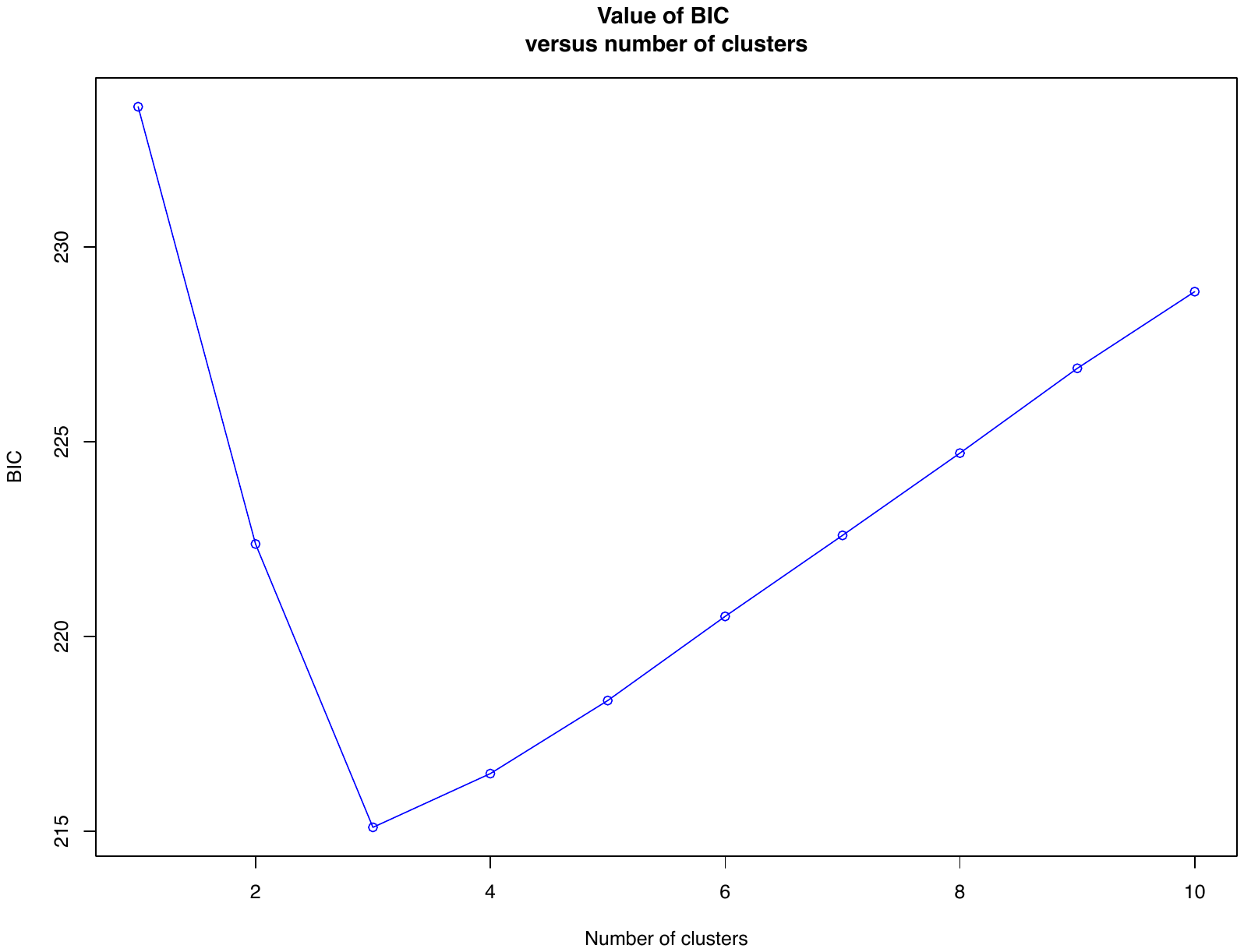

**Figure S19. Results of successive *K*-means clustering for *Bernieria*** (n = 39). The Bayesian Inference Criterion (BIC) is minimized at *K* = 3.

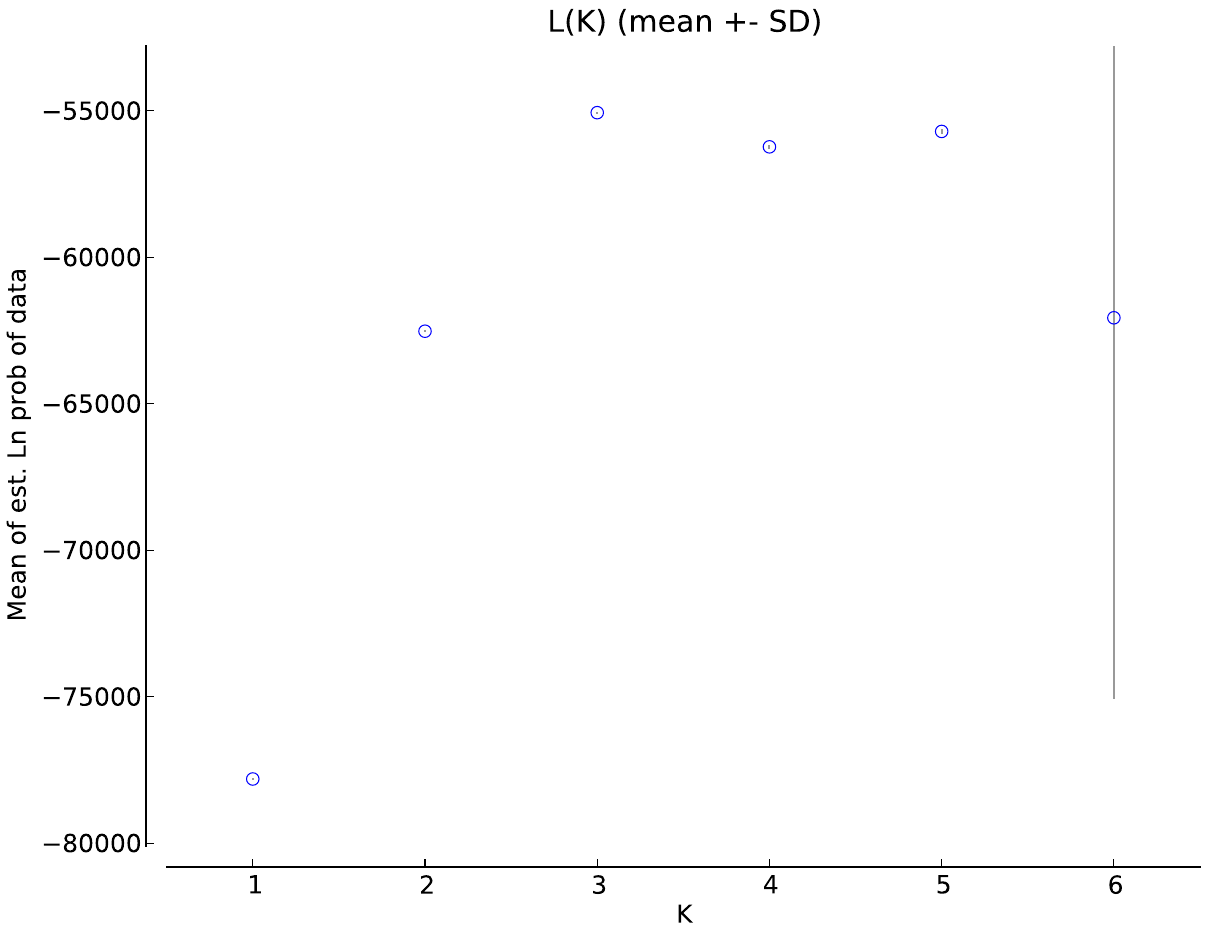

**Figure S20. Estimated log probability of the data Pr(*X*|*K*) for each value of *K* for *Bernieria*** (n = 39), averaged over ten independent Structure runs. The log probability is maximized at *K* = 3.

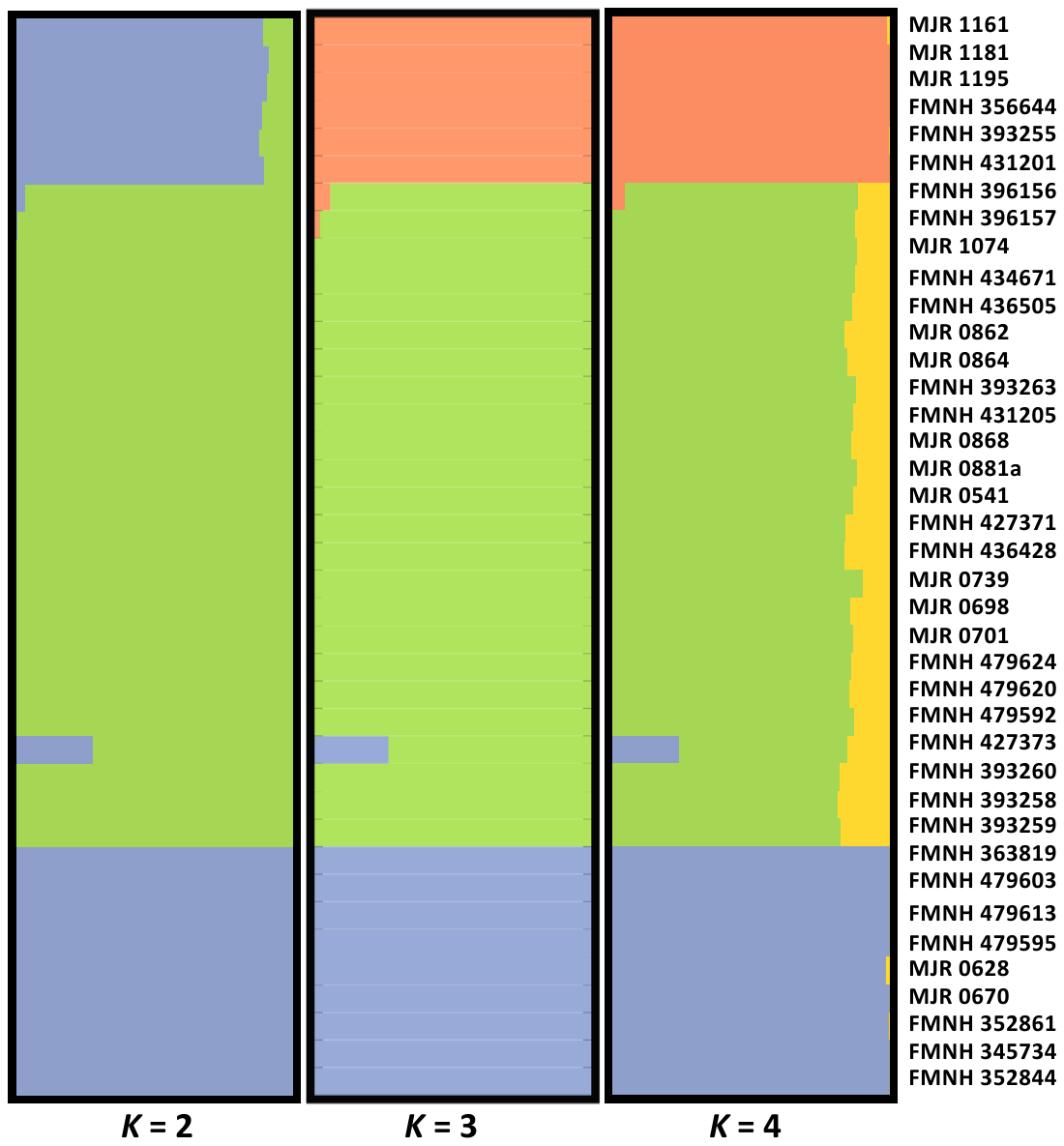

**Figure S21. Structure plots for *Bernieria* (n = 39)**, showing assignment probabilities of each individual to *K* = 2, *K* = 3, and *K* = 4.

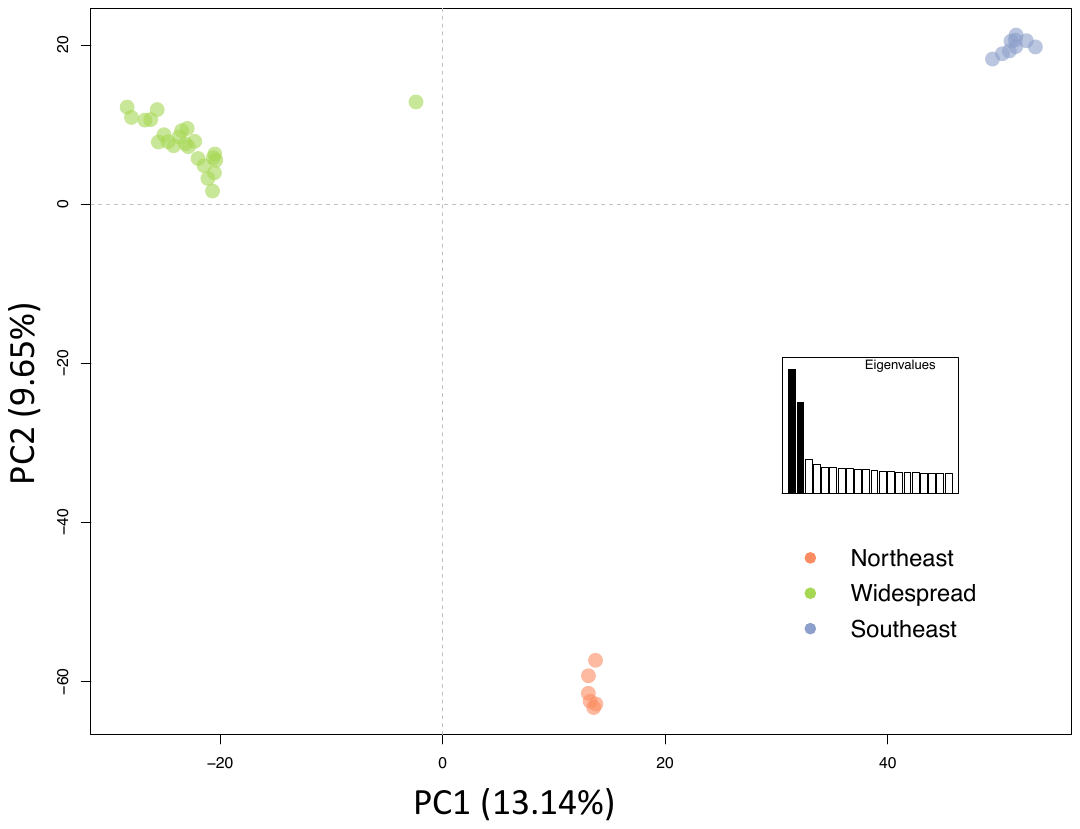

**Figure S22. Principal Components Analysis (PCA) of *Bernieria* (n = 39).**

**Table S3. Pairwise *F_ST_* values between the three *Bernieria* clades.** *F_ST_* values are below the diagonal with *p* values above.

|  | Northeast | Widespread | Southeast |
| --- | --- | --- | --- |
| Northeast | *** | **<0.001** | **<0.001** |
| Widespread | **0.422** | *** | **<0.001** |
| Southeast | **0.439** | **0.454** | *** |

**Table S4. Pairwise *F_ST_* values between the three *Bernieria* clades, with the WS/SE hybrid individual excluded.** *F_ST_* values are below the diagonal with *p* values above.

|  | Northeast | Widespread | Southeast |
| --- | --- | --- | --- |
| Northeast | *** | **<0.001** | **<0.001** |
| Widespread | **0.430** | *** | **<0.001** |
| Southeast | **0.439** | **0.463** | *** |

**Table S5. Morphological measurements (average of three replicates) used in morphological analyses of *Berniera*.** All measurements are in mm. FMNH = Field Museum of Natural History; UADBA = Mention Zoologie Biologie Animale.

| **Accession** | **Clade** | **Sex** | **Bill Length** | **Bill Depth** | **Bill Width** | **Tarsus** | **Wing** | **Tail** | **Simplified Locality** | **Lat** | **Long** |
| --- | --- | --- | --- | --- | --- | --- | --- | --- | --- | --- | --- |
| FMNH 393255 | NE | M | 17.17 | 5.19 | 4.67 | 26.82 | 82.17 | 76.78 | Foret de Betaolana | -14.5 | 49.4 |
| FMNH 393257 | NE | M | 17.46 | 5.45 | 4.83 | 25.75 | 79.78 | 74.35 | Foret de Betaolana | -14.5 | 49.4 |
| UADBA 212145 | NE | F | 13.09 | 4.72 | 4.52 | 24.55 | 73.15 | 75.50 | Maroantsetra | -15.3 | 49.5 |
| UADBA 412127 | NE | F | 12.46 | 4.92 | 4.20 | 24.14 | 69.59 | 70.53 | Maroantsetra | -15.3 | 49.5 |
| UADBA 412146 | NE | F | 13.27 | 4.91 | 4.09 | 25.64 | 69.47 | 69.47 | Maroantsetra | -15.3 | 49.5 |
| UADBA 412147 | NE | F | 13.17 | 4.31 | 4.21 | 24.41 | 71.45 | 68.84 | Maroantsetra | -15.3 | 49.5 |
| UADBA 412149 | NE | F | 13.70 | 4.93 | 4.44 | 24.77 | 70.71 | 72.27 | Maroantsetra | -15.3 | 49.5 |
| UADBA 412151 | NE | F | 12.64 | 4.57 | 4.02 | 24.18 | 72.27 | 66.06 | Maroantsetra | -15.2 | 50.1 |
| UADBA 412153 | NE | F | 13.71 | 5.00 | 4.35 | 23.09 | 72.57 | 64.21 | Maroantsetra | -15.2 | 50.1 |
| UADBA 412156 | NE | F | 13.11 | 5.10 | 4.40 | 23.53 | 72.99 | 71.40 | Maroantsetra | -15.2 | 50.1 |
| UADBA 412129 | NE | M | 15.90 | 5.63 | 4.75 | 24.30 | 79.65 | 70.99 | Maroantsetra | -15.3 | 49.5 |
| UADBA 412133 | NE | M | 15.28 | 5.21 | 4.59 | 27.18 | 78.34 | 73.75 | Maroantsetra | -15.3 | 49.5 |
| UADBA 412134 | NE | M | 17.43 | 5.49 | 4.37 | 26.43 | 85.11 | 80.88 | Maroantsetra | -15.3 | 49.5 |
| UADBA 412137 | NE | M | 16.62 | 5.46 | 4.60 | 25.07 | 84.95 | 76.56 | Maroantsetra | -15.2 | 50.1 |
| UADBA 412138 | NE | M | 16.54 | 5.62 | 4.48 | 23.98 | 83.54 | 77.80 | Maroantsetra | -15.2 | 50.1 |
| UADBA 412140 | NE | M | 16.18 | 5.74 | 4.32 | 26.88 | 84.05 | 76.96 | Maroantsetra | -15.2 | 50.1 |
| UADBA 412142 | NE | M | 17.64 | 5.14 | 3.96 | 25.97 | 88.85 | 82.07 | Maroantsetra | -15.2 | 50.1 |
| UADBA 412143 | NE | M | 16.49 | 5.36 | 4.55 | 26.61 | 82.33 | 72.00 | Maroantsetra | -15.2 | 50.1 |
| UADBA 412144 | NE | M | 16.44 | 5.59 | 4.72 | 24.38 | 85.91 | 83.93 | Maroantsetra | -15.3 | 49.5 |
| FMNH 185918 | SE | F | 12.70 | 5.33 | 4.59 | 25.42 | 70.19 | 65.89 | Manombo | -23 | 47.7 |
| FMNH 345731 | SE | F | 13.39 | 4.82 | 4.66 | 24.96 | 73.42 | 68.37 | Foret de Analalava | -24.2 | 47.3 |
| FMNH 345734 | SE | F | 12.15 | 4.76 | 4.34 | 25.01 | 73.40 | 65.94 | Marosohy Forest | -24.6 | 46.8 |
| FMNH 345740 | SE | F | 12.95 | 4.59 | 4.48 | 24.50 | 74.01 | 63.11 | Marosohy Forest | -24.6 | 46.8 |
| FMNH 345741 | SE | F | 12.20 | 4.80 | 4.70 | 25.48 | 73.32 | 68.60 | Marosohy Forest | -24.6 | 46.8 |
| FMNH 352839 | SE | F | 13.19 | 4.78 | 4.11 | 25.29 | 74.46 | 69.13 | Foret Cascade | -25 | 46.9 |
| **Accession** | **Clade** | **Sex** | **Bill Length** | **Bill Depth** | **Bill Width** | **Tarsus** | **Wing** | **Tail** | **Simplified Locality** | **Lat** | **Long** |
| FMNH 352847 | SE | F | 13.57 | 4.83 | 4.40 | 25.31 | 75.67 | 67.81 | Foret de Marovony | -24.1 | 47.4 |
| FMNH 352860 | SE | F | 13.08 | 4.70 | 4.25 | 25.32 | 70.04 | 65.64 | Foret de Marovony | -24.1 | 47.4 |
| UADBA 412118 | SE | F | 13.91 | 5.36 | 5.23 | 24.75 | 72.71 | 69.32 | Manombo | -23 | 47.7 |
| FMNH 345730 | SE | M | 19.37 | 5.56 | 4.39 | 25.93 | 88.04 | 78.48 | Foret de Analalava | -24.2 | 47.3 |
| FMNH 345732 | SE | M | 16.03 | 5.13 | 4.66 | 25.93 | 83.66 | 74.29 | Marosohy Forest | -24.6 | 46.8 |
| FMNH 345733 | SE | M | 16.70 | 5.79 | 4.56 | 26.52 | 85.17 | 75.60 | Marosohy Forest | -24.6 | 46.8 |
| FMNH 345739 | SE | M | 17.62 | 5.21 | 4.72 | 28.24 | 88.73 | 84.43 | Marosohy Forest | -24.6 | 46.8 |
| FMNH 352838 | SE | M | 17.47 | 5.91 | 5.63 | 27.14 | 88.88 | 81.26 | Foret Cascade | -25 | 46.9 |
| FMNH 352846 | SE | M | 17.75 | 5.51 | 4.54 | 26.56 | 85.62 | 87.65 | Foret de Marovony | -24.1 | 47.4 |
| FMNH 393262 | WS | F | 13.58 | 4.09 | 3.89 | 23.81 | 67.28 | 66.80 | RS d'Ambohijanahary | -18.3 | 45.4 |
| FMNH 396155 | WS | F | 11.93 | 4.18 | 3.75 | 23.68 | 71.45 | 66.82 | Tsinjoarivo | -15.6 | 47.1 |
| FMNH 393259 | WS | M | 17.47 | 5.11 | 4.41 | 26.00 | 85.19 | 89.98 | Manambolo Forest | -22.8 | 47.1 |
| FMNH 393261 | WS | M | 15.94 | 4.81 | 4.11 | 24.92 | 78.69 | 78.52 | RS d'Ambohijanahary | -18.3 | 45.4 |
| UADBA 412177 | WS | F | 12.90 | 4.45 | 4.23 | 22.10 | 73.67 | 73.77 | Tabiky | -22.1 | 44.3 |
| UADBA 412193 | WS | F | 13.58 | 4.57 | 4.53 | 24.51 | 73.39 | 77.31 | Tsiandro | -18.7 | 44.9 |
| UADBA 412195 | WS | F | 12.31 | 4.72 | 4.04 | 23.74 | 69.47 | 74.57 | Tsiandro | -18.7 | 44.9 |
| UADBA 412197 | WS | F | 13.49 | 4.53 | 3.93 | 22.25 | 73.60 | 67.12 | Tsiandro | -18.7 | 44.9 |
| UADBA 412198 | WS | F | 13.32 | 5.06 | 4.46 | 24.82 | 74.15 | 73.23 | Tsiandro | -18.7 | 44.9 |
| FMNH 360044 | WS | M | 14.75 | 4.91 | 4.60 | 24.45 | 77.75 |  | Foret de Zombitsy | -22.9 | 44.7 |
| UADBA 412162 | WS | M | 17.23 | 5.43 | 4.44 | 25.77 | 83.23 | 78.23 | Namoroka | -16.4 | 45.3 |
| UADBA 412194 | WS | M | 17.70 | 4.84 |  | 25.07 | 84.94 | 85.48 | Tsiandro | -18.7 | 44.9 |
| UADBA 412201 | WS | M | 18.91 | 5.98 | 4.55 | 25.22 | 86.40 | 86.70 | Besalampy | -16.8 | 44.9 |
| UADBA 412159 | WS | F | 13.02 | 5.01 | 4.89 | 23.17 | 69.41 | 69.22 | Maromandia | -14.1 | 48.3 |
| UADBA 412160 | WS | F | 12.15 | 4.94 | 4.36 | 24.70 | 72.37 | 70.25 | Maromandia | -14.1 | 48.3 |
| FMNH 185921 | WS | M | 17.63 | 5.30 | 4.22 | 26.91 | 86.45 | 88.24 | Maromandia | -14.1 | 48.3 |
| UADBA 412161 | WS | M | 17.27 | 5.22 | 4.15 | 27.50 | 81.84 | 78.95 | Maromandia | -14.1 | 48.3 |
| FMNH 30967 | WS | M | 16.91 | 5.46 | 4.22 | 26.36 | 81.88 | 73.57 | Forest Sianaka | -18.1 | 48.5 |

**Table S6. Summary statistics of the morphometric measurements taken on museum specimens**. Significant differences (*p* <0.05) between clades in the pairwise ANOVA are indicated by shaded cells.

|  | **Bill Length (BL)** | | **Bill Depth (BD)** | | **Bill Width (BW)** | | **Tarsus** | | **Wing** | | **Tail** | |
| --- | --- | --- | --- | --- | --- | --- | --- | --- | --- | --- | --- | --- |
| **MEANS** | males | females | males | females | males | females | males | females | males | females | males | females |
| NE | 16.65 | 13.14 | 5.44 | 4.81 | 4.53 | 4.28 | 25.76 | 24.29 | 83.15 | 71.53 | 76.92 | 69.79 |
| SE | 17.49 | 13.02 | 5.52 | 4.89 | 4.75 | 4.53 | 26.72 | 25.12 | 86.68 | 73.02 | 80.29 | 67.09 |
| WS | 17.34 | 12.92 | 5.33 | 4.62 | 4.30 | 4.23 | 26.10 | 23.64 | 83.38 | 71.64 | 82.03 | 71.01 |
| **STDEV** |  |  |  |  |  |  |  |  |  |  |  |  |
| NE | 0.73 | 0.44 | 0.20 | 0.26 | 0.24 | 0.18 | 1.15 | 0.77 | 3.11 | 1.47 | 4.10 | 3.54 |
| SE | 1.13 | 0.59 | 0.31 | 0.27 | 0.45 | 0.33 | 0.87 | 0.33 | 2.16 | 1.85 | 5.17 | 2.07 |
| WS | 0.89 | 0.64 | 0.36 | 0.35 | 0.17 | 0.37 | 0.91 | 0.99 | 2.84 | 2.41 | 6.21 | 3.86 |
| **pairwise ANOVA** |  |  |  |  |  |  |  |  |  |  |  |  |
| NE-SE | 0.08 | 0.64 | 0.63 | 0.60 | 0.16 | 0.11 | 0.08 | 0.04 | 0.02 | 0.14 | 0.20 | 0.10 |
| SE-WS | 0.78 | 0.72 | 0.23 | 0.06 | 0.01 | 0.05 | 0.29 | <0.001 | 0.05 | 0.15 | 0.56 | 0.02 |
| NE-WS | 0.12 | 0.42 | 0.38 | 0.19 | 0.10 | 0.69 | 0.49 | 0.09 | 0.86 | 0.92 | 0.05 | 0.45 |

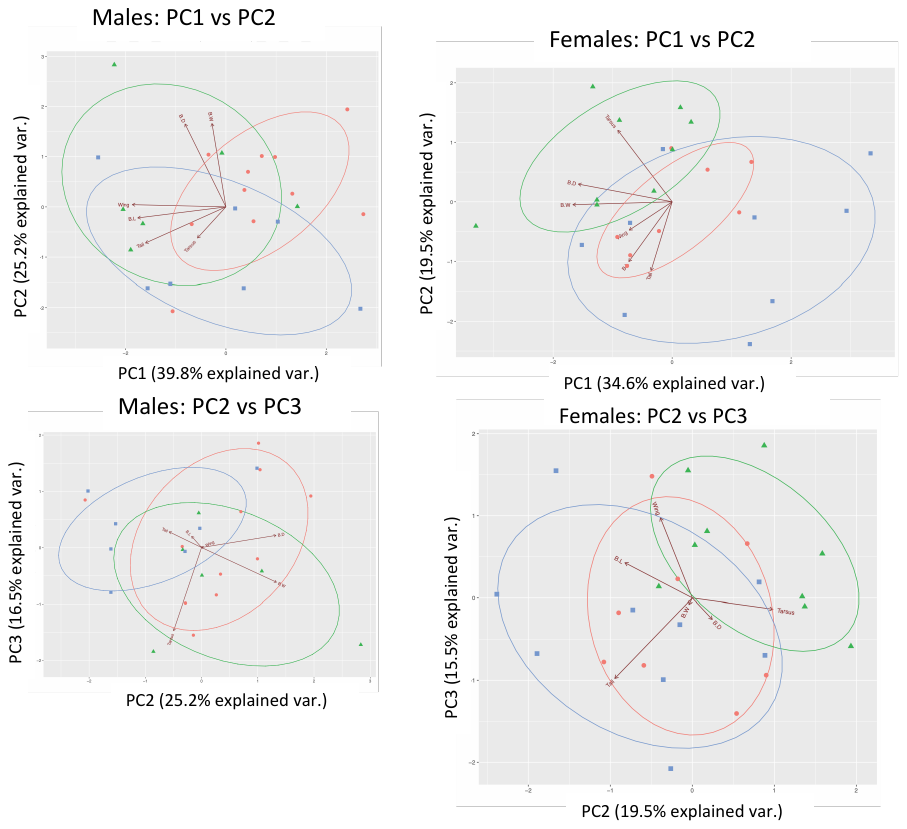

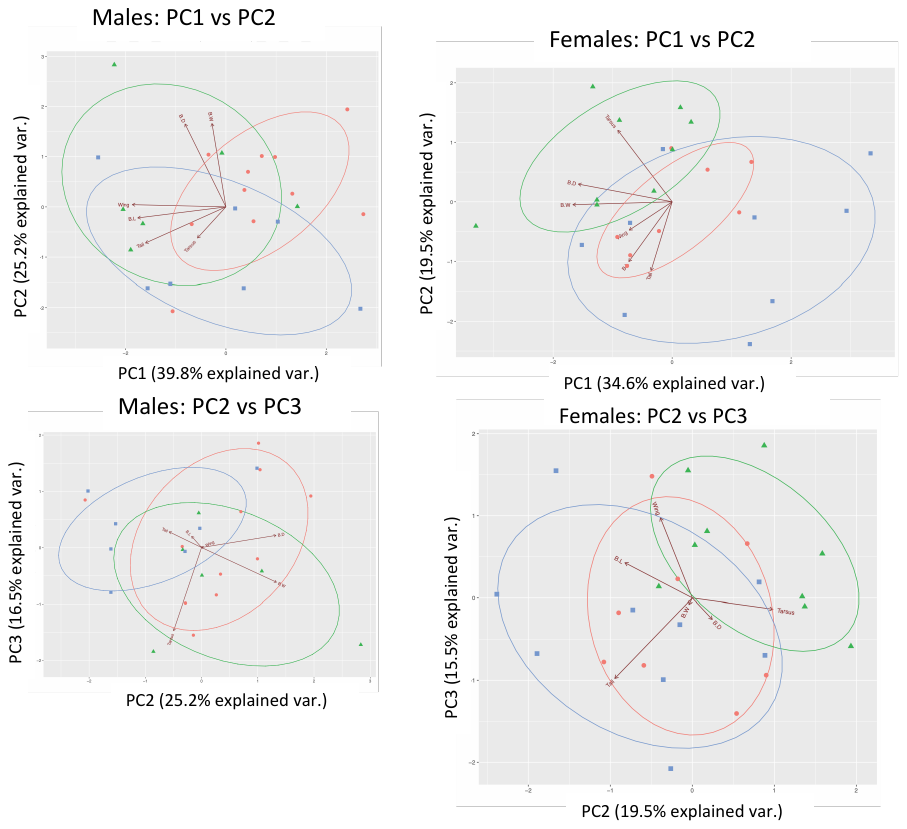

**Figure S23. Principal components analyses (PCAs) of morphological variation in male *Bernieria.*** Males and females were significantly different and were therefore analyzed separately. Centroids of each clade (orange dots = NE; green triangles = SE; blue squares = WS) were significantly different (*p* < 0.001) according to a MANOVA. Circles indicate 95% confidence ellipses around the centroid of each clade.

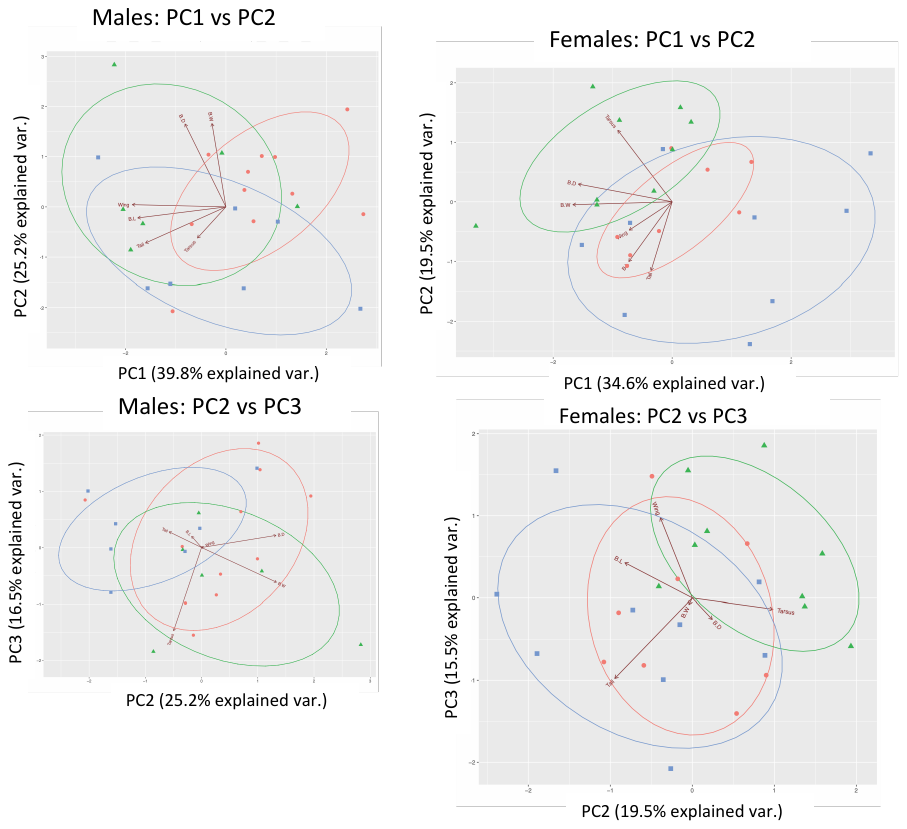

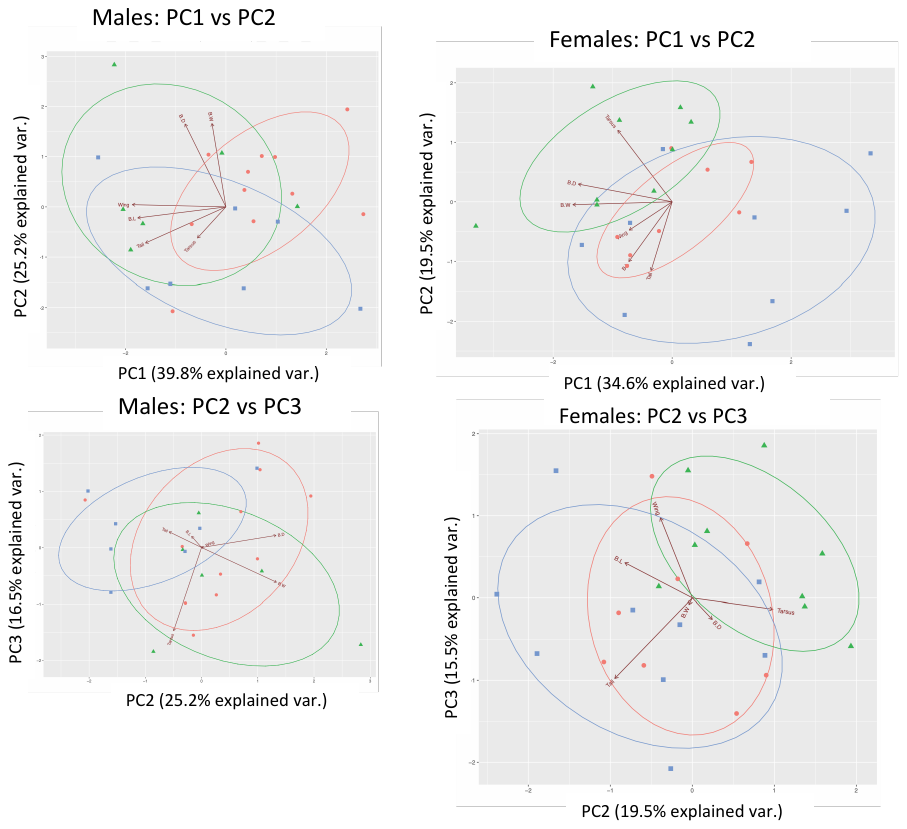

**Figure S24. Principal components analyses (PCAs) of morphological variation in female *Bernieria.*** Males and females were significantly different and were therefore analyzed separately. Centroids of each clade (orange dots = NE; green triangles = SE; blue squares = WS) were significantly different (*p* < 0.001) according to a MANOVA. Circles indicate 95% confidence ellipses around the centroid of each clade.

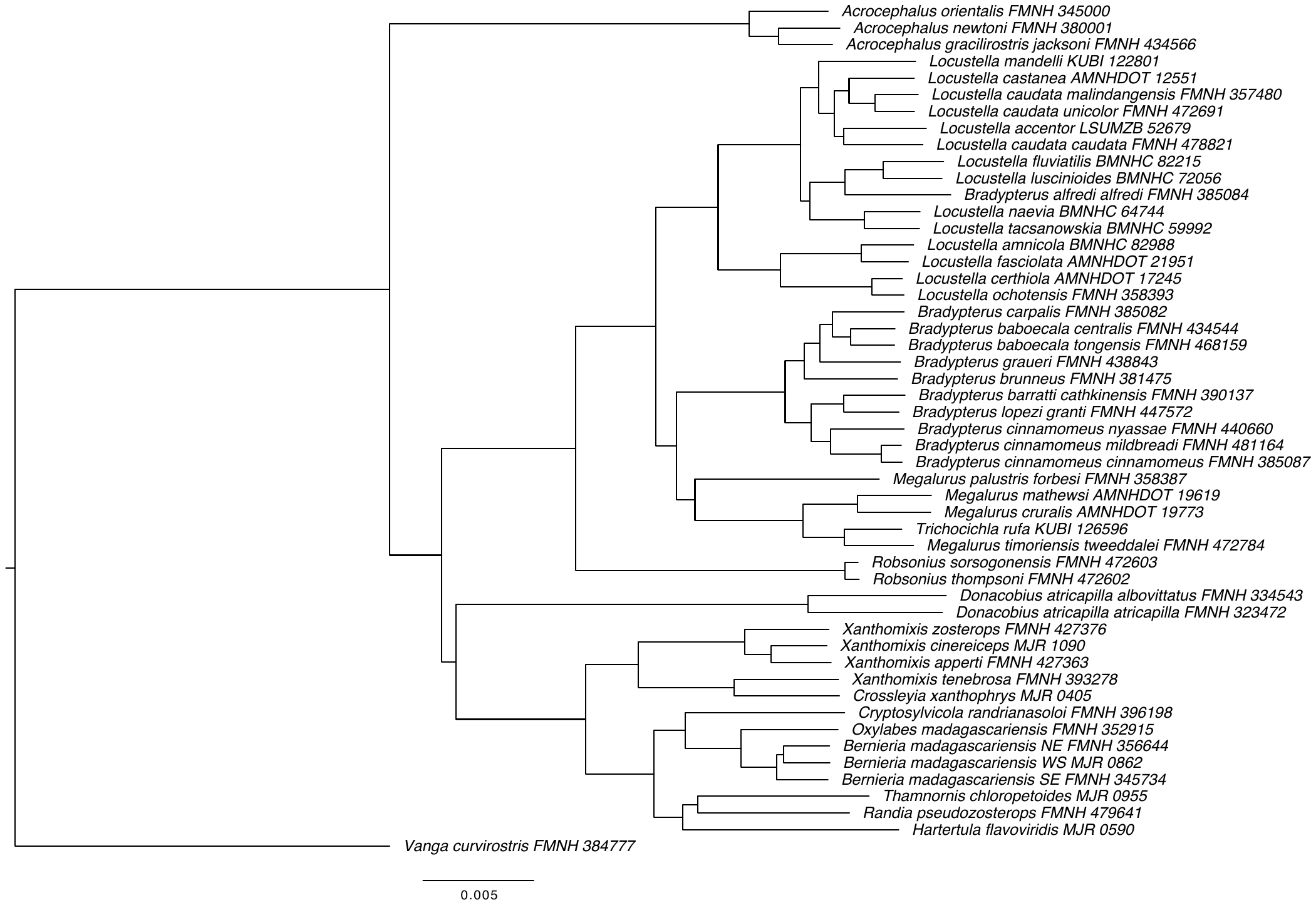

**Figure S25. Maximum-likelihood phylogeny of Bernieridae, Locustellidae, Donacobiidae and Acrocephalidae.** 75% complete data matrix containing 4281 concatenated UCE loci (4,073,553 bp), data were not partitioned. All branches have 100% bootstrap support.

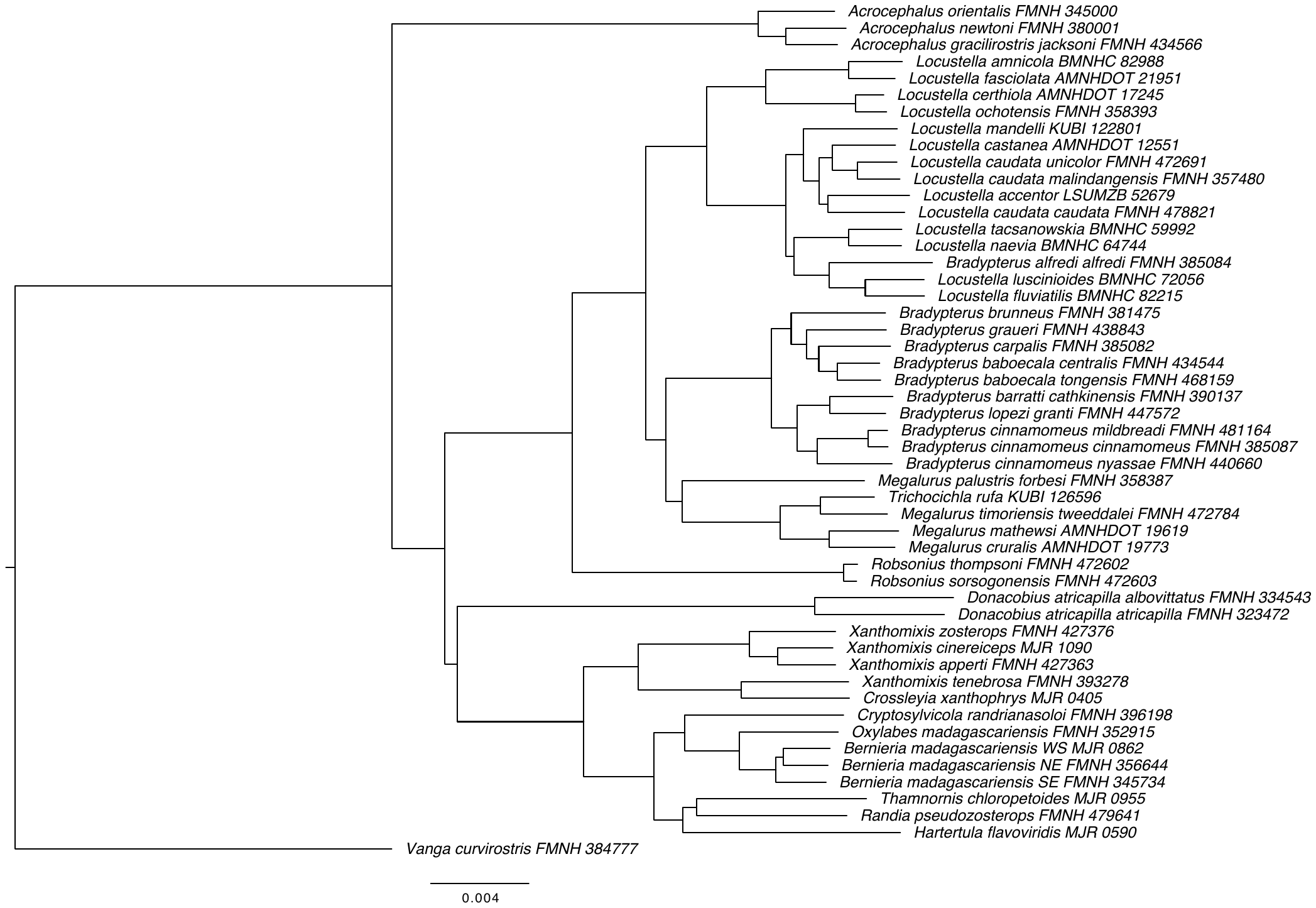

**Figure S26. Maximum-likelihood phylogeny of Bernieridae, Locustellidae, Donacobiidae and Acrocephalidae.** 90% complete data matrix containing 2489 concatenated UCE loci (2,423,537 bp), data were not partitioned. All branches have 100% bootstrap support.

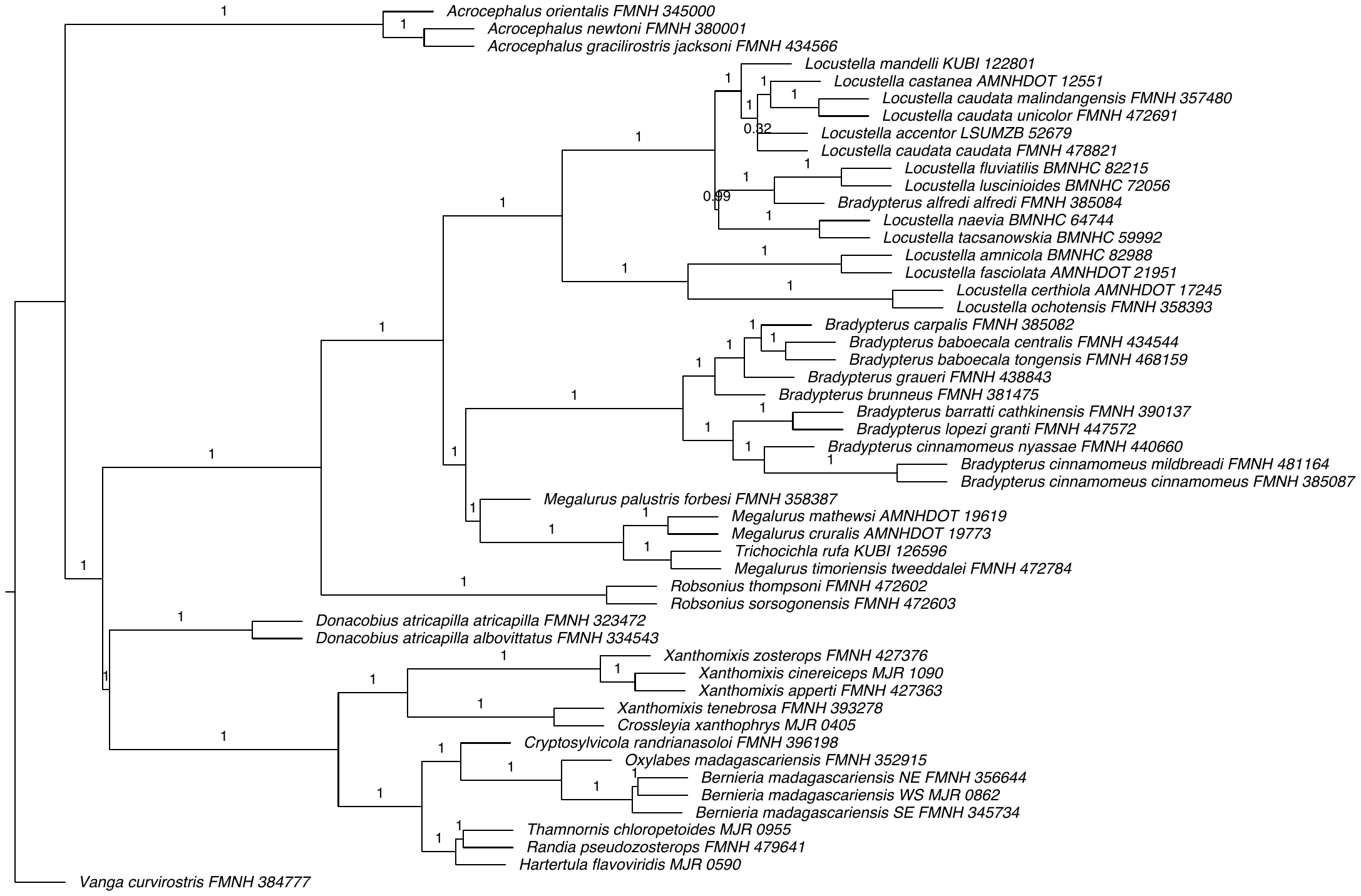

**Figure S27. Species tree of Bernieridae, Locustellidae, Donacobiidae and Acrocephalidae** estimated using ASTRAL from the gene trees of the 1070 most informative UCE loci. Normalized quartet score = 0.93. Branch support values are for quadripartitions (rather than bipartitions). Internal branch lengths are in coalescent units; terminal branch lengths in ASTRAL are arbitrary.
